# Confidence-guided spatial gene expression inference for reliability-aware prognostic analysis from histology

**DOI:** 10.64898/2026.11.23.768521

**Authors:** Xinyi Bai, Renjie Liu, Yuzhi Sun, Guohua Wang, Yadong Wang, Tianyi Zhao

**Affiliations:** Faculty of Computing, Harbin Institute of Technology, Harbin, China; Harbin Institute of Technology Zhengzhou Research Institute, Zhengzhou, China; School of Medicine and Health, Harbin Institute of Technology, Harbin, China

## Abstract

Inferring spatial gene expression from histology images offers a promising strategy for extending spatial molecular profiling to large-scale clinical tissue archives. However, existing approaches face two major challenges: limited generalizability of single-cell expression inference across datasets and platforms, and the lack of reliable estimates of prediction uncertainty, which compromise downstream biological and clinical analyses. Here, we present REGAIN, a framework for spatial gene expression inference at both single-cell and spot resolutions with prediction confidence. Across tissue sections, independent datasets and platforms, REGAIN improved spatial gene expression prediction and downstream analysis. Confidence-guided selection further enriched for cells with more accurately recovered expression profiles and improved cell-type annotation, while confidence-weighted clustering enhanced spatial-domain identification. Beyond assessing prediction quality, confidence mapping reveals greater heterogeneity in gene expression predictability across spatial domains than across broad cell types, with differences persisting within the same cell type and associated with tumor progression-related cell states and local microenvironmental context. Importantly, in 766 patients from the TCGA-BRCA cohort, confidence weighting enhanced the prognostic value of histology-inferred spatial molecular features, yielding stronger prognostic associations and identifying additional prognostic information beyond age and PAM50 molecular subtype. These confidence-weighted features also showed improved prognostic discrimination compared with clinical variables alone or combined with unweighted spatial molecular features. Together, REGAIN marks a shift in histology-based spatial gene expression inference from a focus on predictive accuracy toward a reliability-aware paradigm, demonstrating that prediction confidence provides biological insights into morphology-expression coupling and enhances the clinical informativeness of inferred spatial molecular features.

## Main

Spatial transcriptomics (ST) enables gene expression to be measured while preserving tissue organization, providing a powerful approach for studying intratumoral heterogeneity, cellular states and tumor-microenvironment interactions^1–5^. However, spatial transcriptomic assays remain costly, technically demanding and limited in sample scale^3, 6, 7^. By contrast, hematoxylin and eosin (H&E) staining is routine in pathology, and large collections of digitized H&E slides are available from clinical practice and retrospective cohorts^8–10^. Inferring spatial gene expression from H&E images could therefore extend spatial molecular profiling to large-scale tissue archives.

Histology-based spatial expression prediction has developed rapidly. Early methods mainly focused on spot-level spatial transcriptomics, including iStar^11^, iSCALE^12^, SpaHDmap^13^, scstGCN^14^, STPath^15^, and FmH2ST^16^, which infer or enhance spatial expression from H&E morphology and regional expression constraints. However, spot-level measurements represent averaged signals from multiple cells and therefore cannot directly resolve cellular heterogeneity or fine-scale tissue organization^3, 17^. Recent studies have attempted to extend H&E-to-ST prediction toward cellular resolution, but the level of single-cell supervision differs substantially among existing approaches^18–21^. Methods such as iStar^11^ and iSCALE^12^ generate super-resolved spatial expression maps beyond original sequencing spots but do not provide expression predictions matched to individual cells. sCellST^18^ incorporates cell segmentation to enable cellular expression prediction, yet relies on spot-level aggregate expression supervision rather than paired single-cell expression measurements. Similarly, PRTS^19^ leverages high-resolution Visium HD data to construct finer-grained supervision, although the measured expression units remain spatial bins rather than true single-cell profiles. In contrast, GHIST^20^ and SpatialEx^21^ utilize Xenium-based single-cell spatial transcriptomics to establish matched H&E-single-cell expression pairs, enabling direct cell-level expression prediction from histology. GHIST^20^ established a comprehensive framework for single-cell H&E-to-ST inference by integrating morphology, cell type and neighborhood information, whereas SpatialEx^21^ further focused on extending such predictions across molecular panels and omics modalities.

Despite this progress, most existing single-cell H&E-to-expression models directly learn mappings between histological morphology and transcriptomic profiles, without explicitly modeling the shared biological information between these two modalities. Since morphology and molecular states are intrinsically coupled through cellular identity and tissue organization, learning aligned morphology-transcriptome representations may provide a more transferable basis for single-cell expression inference^22–26^.

Another important limitation is that H&E-derived spatial expression predictions are not uniformly reliable. Because some transcriptional signals are weakly coupled or unrelated to cellular morphology, H&E-derived expression predictions are intrinsically heterogeneous in reliability. Deterministic prediction alone therefore provides insufficient information for downstream use: a predicted expression profile may appear plausible but still be unreliable for cell type annotation or spatial-domain reconstruction^27^. Moreover, deep learning models inherently involve prediction uncertainty due to limitations in data and model representation^28–33^. Therefore, cell-level confidence estimation is necessary to distinguish reliably predictable cells from poorly predictable ones, enabling inferred expression profiles to be used more selectively and accurately in downstream biological and clinical analyses.

To address these challenges, we developed REGAIN, a reliability-aware framework for inferring spatial gene expression from H&E images at single-cell resolution, with adaptability to conventional spot-level prediction. To improve generalizability, REGAIN employs morphology–transcriptome contrastive learning to capture shared biological associations between histological morphology and transcriptional states, rather than relying primarily on dataset-specific visual patterns. Combined with cell–patch joint supervision, which integrates fine-grained cellular information with more robust regional expression constraints, this strategy promotes more transferable morphology–expression representations while reducing sensitivity to noisy cell-level supervision. Importantly, REGAIN explicitly estimates cell-level prediction reliability by learning the relationship between morphological features and empirically measured expression reconstruction errors in held-out cells. This enables REGAIN to infer the expected prediction error of individual cells from their morphology and assign a corresponding confidence score to each predicted expression profile.

We systematically evaluated REGAIN through cross-section, cross-dataset and Xenium-to-Visium HD cross-platform experiments, demonstrating improved single-cell expression recovery and generalizability across diverse tissue sections, independent datasets and spatial transcriptomic platforms. We further validate its applicability to conventional spot-level spatial transcriptomics using the HER2ST dataset, demonstrating its predictive capability across spatial resolutions. We further show that REGAIN outperforms state-of-the-art methods in single-cell expression prediction, while confidence-guided selection further enriches for cells with more accurately recovered expression profiles. Beyond expression recovery, REGAIN achieves superior cell-type annotation and spatial-domain identification, with confidence-guided selection and weighting providing additional improvements in these downstream tasks. We next investigate the biological significance of prediction confidence in breast cancer tissues, revealing how morphology–expression coupling varies across spatial contexts beyond broad cell type identity. Finally, we apply REGAIN to 766 patients from the Cancer Genome Atlas Breast Invasive Carcinoma (TCGA-BRCA) cohort, showing that confidence-informed spatial molecular features provide additional prognostic information beyond conventional prognostic variables and show a trend toward improved survival risk discrimination. Together, we present a generalizable framework for histology-based spatial gene expression inference across resolutions and establish prediction reliability as an important information dimension of molecular inference from histology, extending its value beyond accurate expression reconstruction to biological interpretation and clinical prognostic analysis.

## Results

### Overview of REGAIN

We developed REGAIN, a reliability-aware framework for generalizable single-cell spatial gene expression inference from H&E-stained tissue images (Fig. 1). REGAIN was designed to meet three key requirements for practical histology-based molecular prediction: inferring gene expression at single-cell resolution, generalizing across tissue sections, independent datasets and spatial transcriptomic platforms, and estimating cell-level reliability to distinguish predictable from poorly predictable expression profiles.

**Fig. 1.**
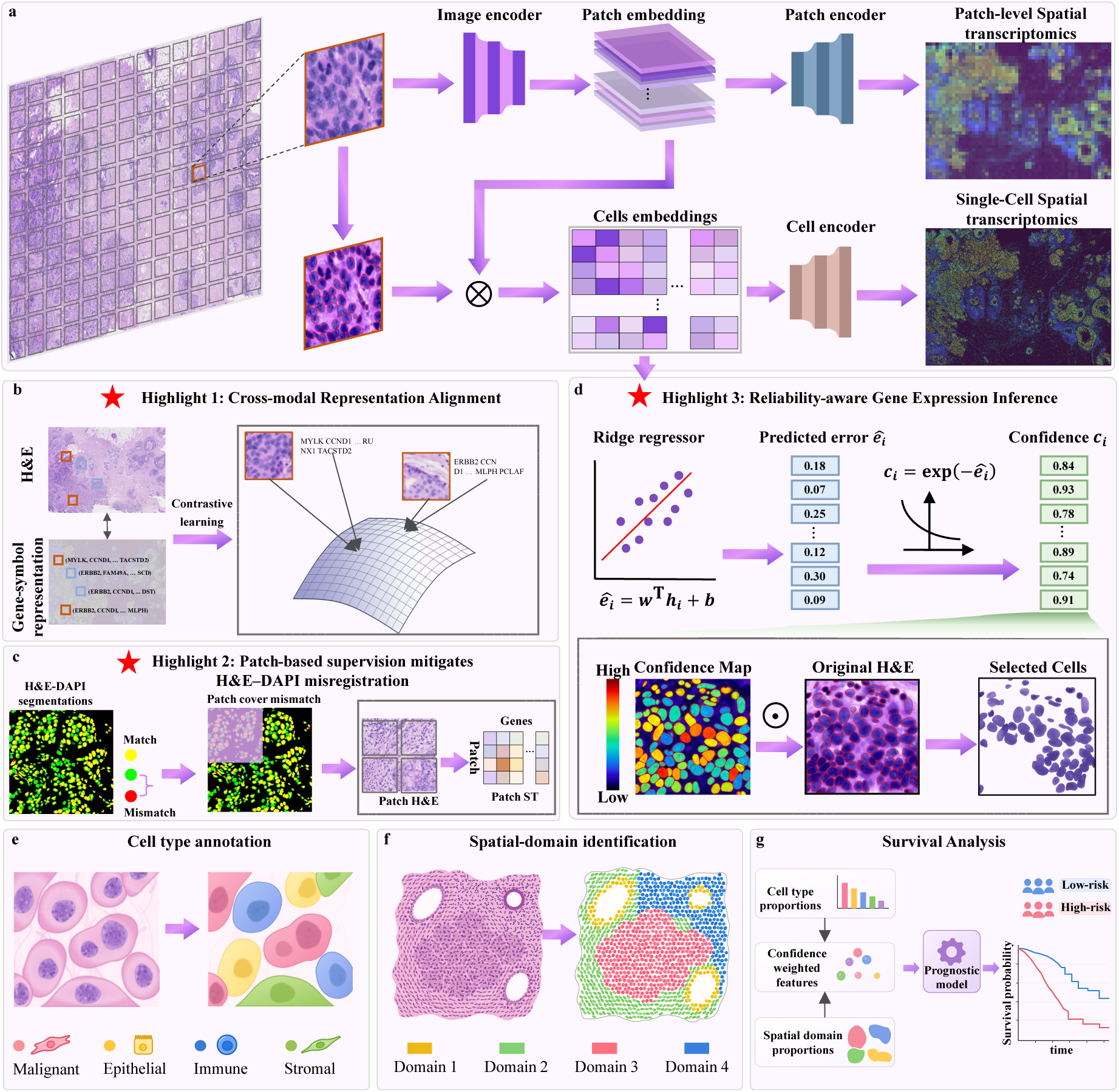
Workflow of REGAIN. (a) Overall workflow for generalizable and reliability-aware single-cell spatial gene expression inference from histology. (b) Morphology-transcriptome contrastive pretraining aligns H&E patch representations with transcriptome-derived gene-symbol representations to learn transcriptionally informative histology features. (c) Cell-patch joint supervision combines single-cell expression labels with patch-level aggregate expression constraints to mitigate the effects of H&E-DAPI registration, segmentation and cell-matching errors. (d) Reliability estimation module, in which a ridge regression model predicts expression error from cell morphology embeddings and converts the predicted error into a confidence score for each cell. (e-g) Downstream applications of REGAIN, including cell type annotation, spatial-domain identification and clinical survival analysis.

REGAIN takes an H&E image and a corresponding cell segmentation mask as input and predicts a spatial gene-expression profile for each detected cell (Fig. 1a). To improve the transferability of morphology-expression representations, REGAIN first performs morphology-transcriptome contrastive pretraining. For each local tissue region, an image encoder extracts histological features from the H&E patch, whereas the paired expression profile is transformed into a transcriptomic semantic representation following the gene-to-text strategy introduced in OmiCLIP^24^, by converting highly expressed genes into gene-symbol sequences and encoding them with a pretrained language encoder. By aligning H&E-derived representations with matched transcriptome-derived representations and separating unmatched pairs (Fig. 1b), this pretraining step encourages the image encoder to capture morphology features associated with local transcriptional states, rather than relying on generic visual patterns alone.

After pretraining, REGAIN is fine-tuned for single-cell expression prediction using both cellular and regional supervision. For each H&E patch, REGAIN extracts cell-level morphological embeddings from segmented cell regions and combines them with the surrounding patch context to predict cell-level expression. In parallel, REGAIN also predicts the aggregate expression of the entire patch from patch-level features (Fig. 1c). This cell-patch joint supervision allows the model to learn from fine-grained single-cell labels while being constrained by more stable regional expression signals. Because single-cell training labels depend on cell segmentation and H&E-DAPI registration and matching, patch-level supervision provides an additional regional constraint that reduces the model’s sensitivity to local cell-boundary and cell-matching errors.

A key component of REGAIN is its cell-level reliability estimation. During model development, held-out training cells are used to estimate empirical prediction errors. REGAIN then learns the relationship between cell morphological embeddings and expression reconstruction errors using a ridge regression model. During inference on a new section, the ridge model estimates the expected prediction error for each cell, and converts it into a confidence score (Fig. 1d). Thus, REGAIN outputs not only a predicted single-cell expression profile but also a cell-level confidence score that reflects the expected quality of the prediction.

The predicted expression profiles can be used for downstream spatial transcriptomic analyses, including cell type annotation, spatial-domain identification and clinical survival analysis (Fig. 1e-g). Importantly, prediction confidence can be incorporated into these analyses through confidence-guided selection and weighting, improving downstream analytical performance and enhancing the prognostic value of inferred spatial molecular features. Together, REGAIN integrates generalizable single-cell expression inference with reliability-aware quality assessment, enabling quality-assessed spatial molecular profiling from routine histology.

### REGAIN enables confidence-guided cross-section inference of single-cell spatial expression

We first evaluated whether REGAIN could generalize from one tissue section to another section in a breast cancer Xenium dataset^34^. REGAIN was trained on one section (BreastCancer1 Rep1) and tested on another section (BreastCancer1 Rep2), providing a stringent cross-section setting in which the model had to infer single-cell spatial expression in an unseen tissue section from H&E morphology alone.

After H&E-based cell segmentation and matching to Xenium-derived expression profiles, 127,964 matched cells were obtained from the training section. We used 24,666 cells to train the reliability estimator and the remaining cells to train the main expression prediction model. The rep2 section contained 79,168 matched cells and was reserved for testing. The Xenium panel contained 313 genes, and the median segmented cell area was approximately 25 μm². Details of cell segmentation, image registration and cell matching are provided in the Methods.

REGAIN improved gene-expression recovery compared with GHIST^20^ across all evaluated gene sets, including all genes, highly variable genes (HVGs) and spatially variable genes (SVGs) (Fig. 2a). These results indicate that REGAIN better preserves both overall expression variation and spatially structured transcriptional signals in an unseen adjacent section.

**Fig. 2.**
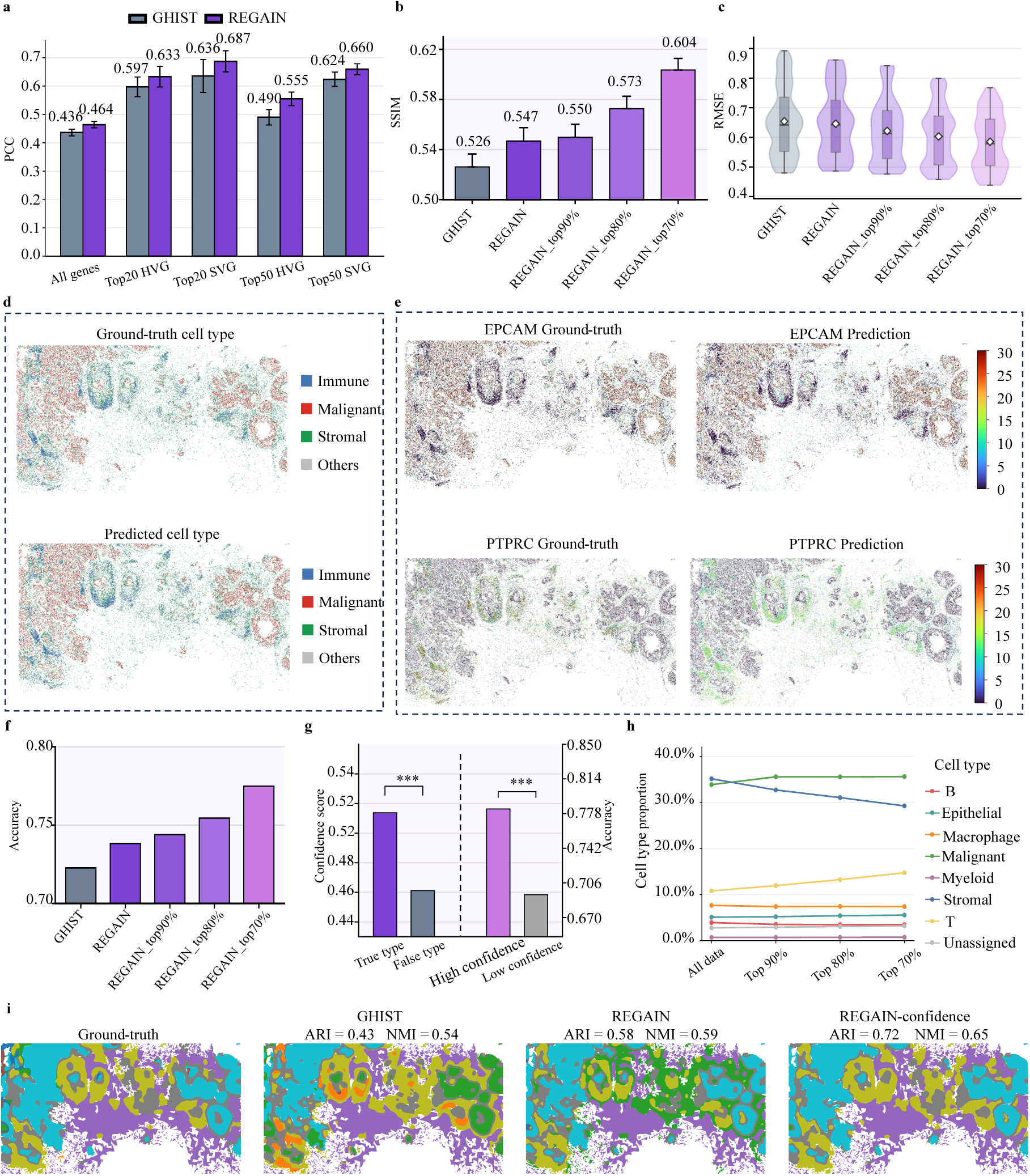
Confidence-guided cross-section inference on consecutive Xenium breast cancer sections. (a) Gene-wise Pearson correlation coefficient (PCC) comparison between GHIST^20^ and REGAIN for all genes, HVGs and SVGs. (b) Gene-wise SSIM comparison for GHIST^20^, REGAIN and REGAIN predictions after retaining the top 90%, 80% and 70% most confident cells. (c) Cell-wise expression RMSE comparison for GHIST^20^, REGAIN and confidence-filtered REGAIN predictions. (d) Ground-truth and predicted cell type maps using a broad marker-associated grouping, including immune, malignant, stromal and other cells. (e) Spatial visualization of representative marker genes. EPCAM was used as a malignant epithelial/tumor-cell marker, and PTPRC was used as an immune-cell marker. For each gene, ground-truth expression is shown in the left column and REGAIN-predicted expression in the right column. (f) Cell type annotation accuracy based on the GHIST-style cell type grouping. Results are shown for GHIST^20^, REGAIN and REGAIN predictions after confidence-guided filtering. (g) Relationship between confidence and cell type annotation correctness. The left panel compares confidence scores between correctly and incorrectly annotated cells, and the right panel compares annotation accuracy between high- and low-confidence cells. Statistical significance was assessed between the two bars in each comparison; *** indicates P < 0.001. (h) Changes in cell type proportions after confidence-guided filtering, based on the conventional GHIST-style cell type grouping, including B, Epithelial, Macrophage, Malignant, Myeloid, Stromal, T and Unassigned cells. (i) Spatial-domain identification based on measured expression, GHIST-predicted expression, REGAIN-predicted expression and REGAIN-predicted expression incorporating cell-level prediction confidence. ARI and NMI were computed by comparing the spatial domains identified by each prediction-based approach with those inferred from measured expression.

Beyond improving overall prediction accuracy, REGAIN also assigns a confidence score to each predicted cell, allowing the predicted expression map to be stratified by estimated reliability. We next assessed whether the confidence score could identify cell-level predictions that are more suitable for downstream analysis. Confidence-guided filtering progressively improved spatial expression recovery, with the top 90%, 80% and 70% most confident cells showing increased gene-wise structural similarity index measure (SSIM) and reduced cell-wise root mean squared error (RMSE) compared with unfiltered predictions (Fig. 2b, c). Thus, confidence scores provided a practical reliability layer for selecting higher-quality inferred expression profiles.

REGAIN also recovered biologically coherent marker-gene patterns. In the test section, cell types were grouped into malignant, immune, stromal and other cells to enable direct comparison with marker-gene expression patterns (Fig. 2d). Cell-type annotation based on REGAIN-predicted expression achieved an accuracy of 0.81 under this four-category grouping. REGAIN recovered spatially coherent expression patterns for representative marker genes, including EPCAM for malignant epithelial/tumor cells and PTPRC for immune cells (Fig. 2e). The predicted marker-gene distributions broadly corresponded to the spatial organization of the associated cell types, supporting the biological consistency of the inferred expression. Similar trends were also observed for the stromal marker LUM, whose predicted expression aligned with stromal-enriched regions (Supplementary Fig. 1a). Additional breast cancer- or tumor-associated marker genes also showed consistent spatial recovery patterns (Supplementary Fig. 1b). These results support the biological consistency of the inferred single-cell expression profiles.

We next evaluated whether REGAIN-predicted expression profiles could improve downstream cell-type annotation. We applied the neural-network classifier described in Methods to the gene-expression profiles predicted by REGAIN and GHIST^20^, respectively. The fine-grained cell-type annotations provided by the original dataset authors were merged into eight broader categories following the grouping scheme used in GHIST^20^: macrophages, malignant cells, stromal cells, T cells, epithelial cells, B cells, myeloid cells and unassigned cells. These grouped annotations were used as reference labels to evaluate cell-type annotation accuracy. Cell type annotation based on REGAIN-predicted expression was more accurate than that based on GHIST^20^ predictions, and confidence-guided filtering further increased annotation accuracy (Fig. 2f). Correctly annotated cells showed higher confidence scores than incorrectly annotated cells, and high-confidence cells achieved higher annotation accuracy than low-confidence cells (Fig. 2g). Confidence-guided filtering further altered the inferred cell type composition, with a reduced proportion of stromal cells and an increased proportion of malignant cells as the confidence threshold became more stringent (Fig. 2h, Supplementary Fig. 2). This pattern may reflect differences in morphology–expression predictability across cellular populations, with malignant cells potentially exhibiting more readily recognizable morphology– expression associations, whereas transcriptional states in some stromal populations may be less reliably inferred from H&E morphology alone.

We next evaluated whether REGAIN-predicted expression could support spatial-domain identification and whether incorporating prediction confidence could further improve its performance. Rather than filtering out low-confidence cells, we incorporated cell-level confidence scores as weights in KMeans clustering, allowing all cells to contribute to spatial-domain identification while assigning greater influence to cells with more reliable predicted expression profiles. Notably, the spatial distribution of prediction confidence exhibited distinct regional patterns (Supplementary Fig. 3), which were further leveraged to refine the initial clustering results and improve spatial-domain identification (Details see Methods). Compared with GHIST^20^, spatial domains inferred from REGAIN-predicted expression showed greater agreement with those derived from measured expression, achieving higher ARI (0.58 vs. 0.43) and NMI (0.59 vs. 0.54) (Fig. 2i). Incorporating prediction confidence further improved spatial-domain identification(see methods), yielding the highest ARI (0.72) and NMI (0.65) among the evaluated methods. Consistent with these quantitative improvements, the confidence-guided REGAIN results showed the closest visual correspondence to the ground-truth spatial domains, better preserving the spatial organization and boundaries of tissue compartments (Fig. 2i).

Together, these results demonstrate that REGAIN enables accurate single-cell spatial gene expression prediction across tissue sections and supports reliable downstream cell-type annotation and spatial-domain identification. Importantly, confidence-guided selection not only further improves expression recovery among retained cells, but also enables more reliable downstream analysis.

### REGAIN generalizes confidence-guided single-cell spatial expression inference across independent datasets

We next evaluated REGAIN in a more challenging cross-dataset setting, in which the model was trained on one Xenium section and tested on a section from an independent dataset. Compared with the adjacent-section experiment, this setting introduced additional variation in tissue composition, staining appearance and cellular organization.

In this experiment, the training data were derived from the 10x Genomics FFPE human breast dataset. The source dataset (BreastCancer2) contained 114,996 matched cells measured with a 280-gene Xenium panel. We used 16,554 cells to train the reliability estimator. The remaining 98,442 cells were used to train the main gene-expression prediction model. The independent TENX94 Xenium breast cancer dataset was used as the cross-dataset test sample. It shared the same 280-gene panel and contained 275,628 matched cells. The cell segmentation, cell matching, cell-area distribution and cell type grouping followed the same procedure as in the cross-section experiment.

REGAIN maintained improved expression recovery over GHIST^20^ across all evaluated gene sets, including all genes, HVGs and SVGs (Fig. 3a). Confidence-guided filtering further improved spatial pattern preservation and reduced cell-wise prediction error, as shown by increased gene-wise SSIM and decreased RMSE among high-confidence cells (Fig. 3b, c).

**Fig. 3.**
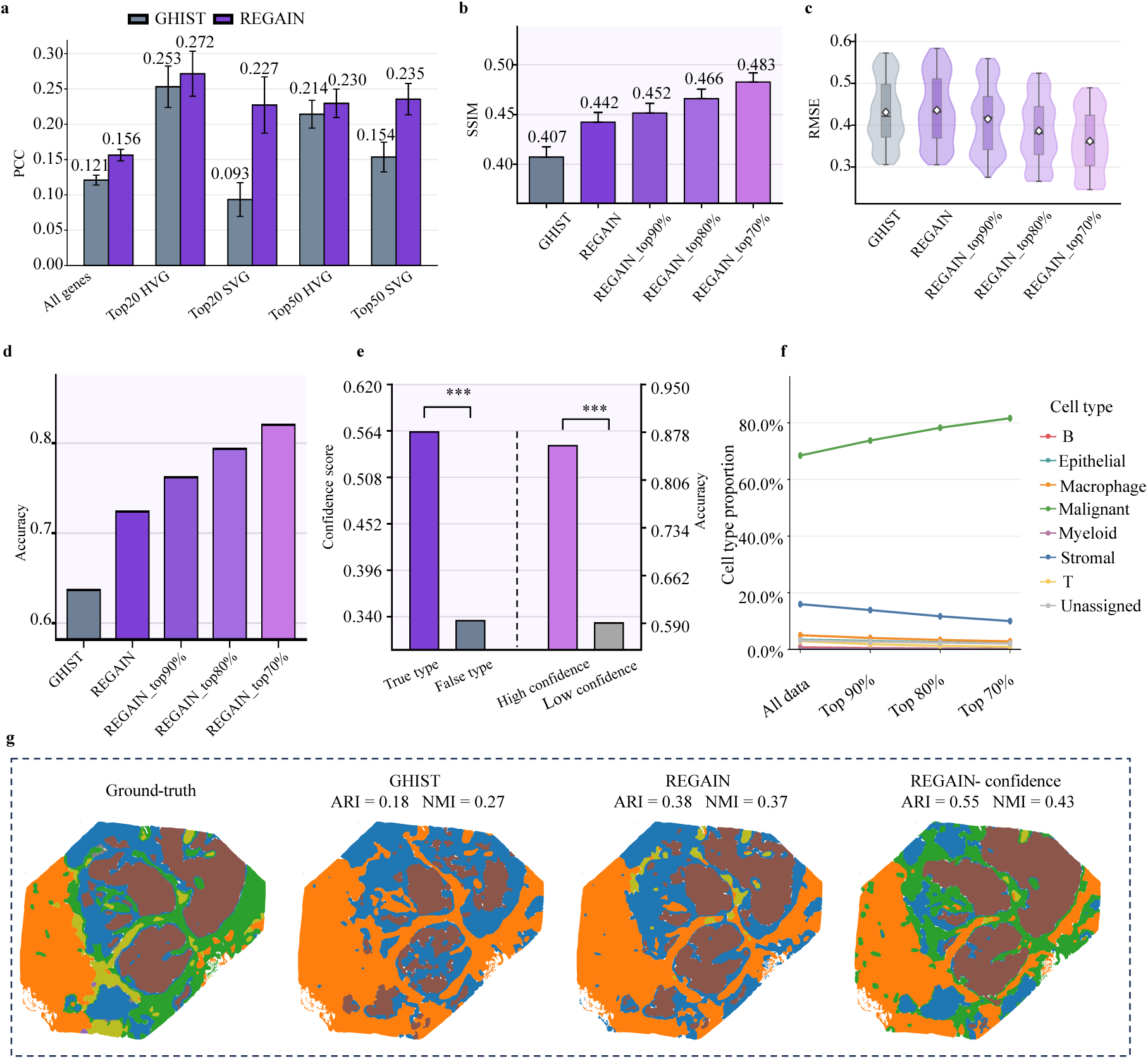
Confidence-guided cross-dataset inference on independent Xenium breast cancer sections. (a) Gene-wise PCC comparison between GHIST^20^ and REGAIN across all genes, the top 20 HVGs, the top 20 SVGs, the top 50 HVGs and the top 50 SVGs. (b) Gene-wise SSIM comparison for GHIST^20^, REGAIN and REGAIN predictions after confidence-guided filtering. (c) Cell-wise expression RMSE comparison for GHIST^20^, REGAIN and confidence-filtered REGAIN predictions. (d) Cell-type annotation accuracy based on the conventional GHIST-style cell-type grouping. Results are shown for GHIST^20^, REGAIN and REGAIN predictions after confidence-guided filtering. (e) Relationship between prediction confidence and cell-type annotation correctness. The left panel compares confidence scores between correctly and incorrectly annotated cells, and the right panel compares annotation accuracy between high- and low-confidence cells. Statistical significance was assessed between the two bars in each comparison; *** indicates P < 0.001. (f) Changes in cell-type proportions across confidence-filtering thresholds, comparing all cells with subsets retaining the top 90%, 80% and 70% most confident cells. (g) Spatial-domain identification based on measured expression, GHIST-predicted expression, REGAIN-predicted expression and confidence-guided REGAIN predictions. ARI and NMI were computed by comparing the spatial domains identified by each prediction-based approach with those inferred from measured expression.

We further examined whether REGAIN-predicted expression could recover representative cellular compartments, including immune-, malignant- and stromal-enriched regions, in the cross-dataset setting. The predicted cell-type distribution showed overall consistency with the ground-truth cell-type map, achieving an annotation accuracy of 0.74 under the four-category grouping (Supplementary Fig. 4). We then examined representative marker-gene expression patterns to further assess the biological consistency of the inferred expression profiles. REGAIN recovered spatially coherent expression patterns for EPCAM, PTPRC and LUM, corresponding to malignant epithelial/tumor cells, immune cells and stromal cells, respectively. The predicted expression patterns broadly matched those derived from measured expression and aligned with the spatial distribution of their associated cellular compartments (Supplementary Fig. 5). Additional breast cancer- or tumor-associated marker genes, including ERBB2 and ESR1, also showed consistent spatial recovery patterns (Supplementary Fig. 6).

Consistent improvements were also observed for expression-derived cell type annotation in the cross-dataset setting. As the TENX94 dataset did not provide cell-type annotations, annotations derived from measured gene expression were used as reference labels, following the same eight-category grouping scheme as in the cross-section analysis. REGAIN achieved higher annotation accuracy than GHIST^20^ (0.72 versus 0.64), and retaining the top 70% high-confidence cells further improved the accuracy to 0.82 (Fig. 3d). Correctly annotated cells showed a higher mean confidence score than incorrectly annotated cells (0.56 versus 0.34), and high-confidence cells achieved higher annotation accuracy than low-confidence cells (0.86 versus 0.59; Fig. 3e). Confidence-guided filtering also altered the composition of annotated cell types, with a decreased proportion of stromal cells and an increased proportion of malignant cells under more stringent confidence selection (Fig. 3f, Supplementary Fig. 7). This trend was consistent with the cross-section analysis, suggesting reproducible, cell-population-associated differences in prediction confidence across datasets.

Finally, we evaluated whether REGAIN-predicted expression could support spatial-domain identification in the cross-dataset setting and whether incorporating prediction confidence could further improve its performance. Following the same strategy used in the cross-section analysis, spatial domains were identified from the processed spatial gene expression matrices using KMeans clustering. Cell-level prediction confidence was incorporated into the clustering process, and its spatial distribution, which exhibited distinct regional patterns (Supplementary Fig. 8), was further leveraged to refine the initial clustering results (see Methods for details). Compared with GHIST^20^, spatial domains inferred from REGAIN-predicted expression showed greater agreement with those derived from measured expression, achieving higher adjusted Rand index (ARI; 0.38 vs. 0.18) and normalized mutual information (NMI; 0.37 vs. 0.27) (Fig. 3g). Incorporating prediction confidence further improved spatial-domain identification, increasing ARI from 0.38 to 0.55 and NMI from 0.37 to 0.43. Consistent with these quantitative improvements, confidence-guided REGAIN produced spatial-domain patterns that more closely resembled the ground-truth domains, better preserving the spatial organization of tissue compartments (Fig. 3g).

Together, these results demonstrate that REGAIN generalizes across independent datasets, while its prediction confidence helps identify unreliable expression predictions and improves the performance of downstream analyses. We further evaluated cross-platform transfer from Xenium to Visium HD^35^ in breast and lung cancer, demonstrating the applicability of REGAIN to different spatial transcriptomic platforms with limited target-platform adaptation (Supplementary Note 1 and Supplementary Fig. 9).

### REGAIN confidence reveals regional and cell-type identity heterogeneity in morphology-associated predictability

We next sought to understand whether morphology–expression predictability is primarily associated with cellular identity or shaped by the local tissue microenvironment. Unlike conventional evaluation metrics that summarize overall prediction performance, cell-level confidence enables the reliability of individual inferred expression profiles to be examined in their spatial and cellular contexts. This analysis therefore aimed to determine whether REGAIN confidence could identify which molecular profiles are more reliably predictable from H&E morphology beyond expression accuracy alone.

To investigate this question, we performed reciprocal cross-section prediction using two consecutive breast cancer Xenium sections, BreastCancer1 Rep1 and BreastCancer1 Rep2. Specifically, REGAIN was first trained on BreastCancer1 Rep1 and evaluated on BreastCancer1 Rep2, and the training and test sections were then reversed. This experimental design enabled us to examine morphology– expression predictability in both tissue sections while maintaining independent training and testing. We analyzed cell-level prediction confidence and prediction accuracy across cell types and spatial domains, and further examined the similarity of their H&E-derived morphological representations. Unless otherwise specified, confidence and prediction accuracy distributions were pooled from the two cross-section experiments.

We first examined how morphology–expression predictability varied across cell types. Cell-level confidence scores exhibited substantial heterogeneity among different cellular populations, with malignant cells showing the highest confidence, whereas several other cell types exhibited lower and more variable confidence distributions (Fig. 4a). A similar pattern was observed when directly evaluating expression prediction accuracy using cell-wise PCCs between predicted and measured expression profiles across all 313 genes (Fig. 4b). Malignant cells achieved the highest PCCs, broadly consistent with their elevated prediction confidence. To explore whether these differences were associated with morphological distinguishability, we further compared H&E-derived morphological embeddings across cell-type groups. The resulting similarity matrices revealed distinct patterns of morphological similarity among cell types, with malignant cells exhibiting more distinctive morphological features that differed from those of other cellular populations. Similar patterns were observed in both BreastCancer1 Rep1 and BreastCancer1 Rep2 (Fig. 4c). Together, these observations suggest that differences in morphology–expression predictability across cell types may be associated, at least in part, with the distinctiveness of their histological features.

**Fig. 4.**
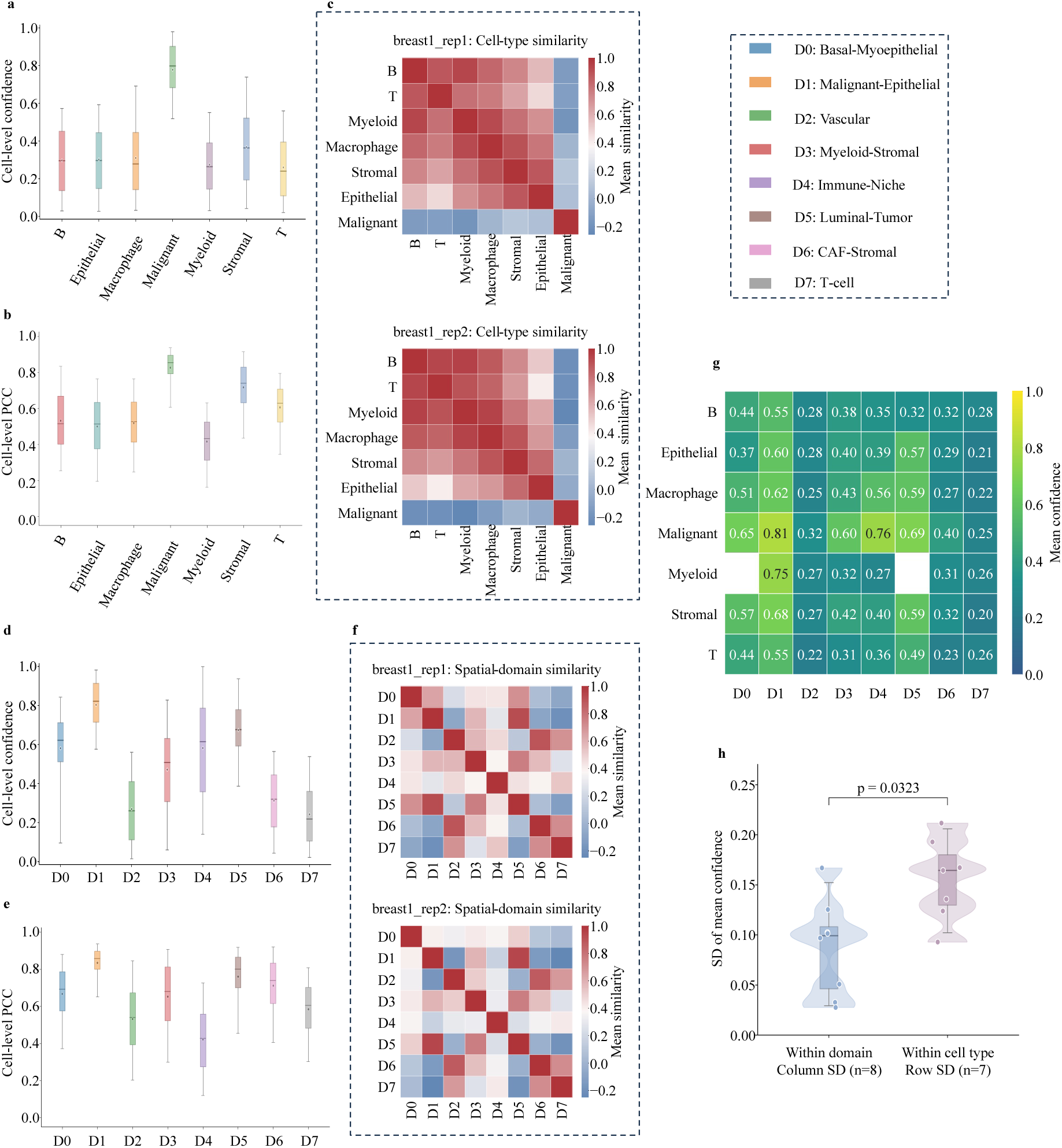
REGAIN confidence reveals regional and cell-type heterogeneity in morphology-associated predictability. Analyses were performed on two consecutive breast cancer Xenium sections (BreastCancer1 Rep1 and BreastCancer1 Rep2) using reciprocal cross-section prediction, with each section serving alternately as the training and test section. (a) Distribution of cell-level prediction confidence across cell types, pooled from the two cross-section experiments. (b) Distribution of cell-wise PCCs between predicted and measured gene expression profiles across cell types, pooled from the two cross-section experiments. (c) Heatmaps showing pairwise similarities between cell-type groups in the H&E-derived morphological embedding space, evaluated separately for BreastCancer1 Rep1 (top) and BreastCancer1 Rep2 (bottom). (d) Distribution of cell-level prediction confidence across eight spatial domains, pooled from the two cross-section experiments. (e) Distribution of cell-wise PCCs between predicted and measured expression profiles across spatial domains, pooled from the two cross-section experiments. (f) Heatmaps showing pairwise similarities between spatial-domain groups in the morphological embedding space, evaluated separately for BreastCancer1 Rep1 (top) and BreastCancer1 Rep2 (bottom). (g) Heatmap showing the mean prediction confidence for cells belonging to each cell-type–spatial-domain combination. Rows represent cell types, and columns represent spatial domains. (h) Comparison of confidence variability within spatial domains and within cell types, quantified as the standard deviation (SD) of the mean confidence values across cell types within each spatial domain (columns, n = 8) and across spatial domains within each cell type (rows, n = 7). Statistical significance was assessed between the two groups (P = 0.0323).

We next investigated whether similar heterogeneity existed across spatial domains. The two sections were jointly integrated using STAligner^36^ to identify shared spatial domains (see Methods for details). These domains were subsequently annotated based on their enriched cell-type composition and domain-specific marker genes, yielding eight spatial domains: basal-myoepithelial (D0), malignant-epithelial (D1), vascular (D2), myeloid-stromal (D3), immune-niche (D4), luminal-tumor (D5), cancer-associated fibroblast stromal (CAF-stromal; D6) and T-cell-enriched (D7). Prediction confidence varied markedly across these spatial domains (Fig. 4d). In particular, the malignant-epithelial domain (D1) exhibited relatively high confidence, whereas the vascular domain (D2) and T-cell-enriched domain (D7) showed lower confidence levels. Cell-wise PCCs also differed substantially among spatial domains, with the malignant-epithelial domain achieving the highest prediction accuracy and the vascular domain showing comparatively poor expression recovery (Fig. 4e). We further examined morphological similarities among spatial domains using H&E-derived embeddings. The resulting similarity matrices revealed distinctive morphological characteristics across spatial domains in both sections (Fig. 4f). Together, these results demonstrate that morphology–expression predictability is heterogeneous across both cellular populations and spatial domains, with distinct patterns of prediction confidence, expression recovery accuracy and morphological characteristics across different tissue contexts.

To further disentangle the associations of cell identity and spatial context with prediction confidence, we jointly examined confidence scores across cell types and spatial domains. For each cell-type–spatial-domain combination, we calculated the mean prediction confidence of the corresponding cells (Fig. 4g). The resulting heatmap revealed that confidence scores were relatively consistent across different cell types within the same spatial domain, whereas substantial variation was observed across spatial domains within individual cell types. For example, malignant cells exhibited markedly different confidence levels across spatial domains, despite sharing the same cell-type identity. To quantify this pattern, we calculated the standard deviation of mean confidence values across cell types within each spatial domain and across spatial domains within each cell type. Variability within spatial domains was significantly lower than variability within cell types (P = 0.0323; Fig. 4h), indicating that morphology– expression predictability is more strongly associated with local spatial context than with broad cell-type identity.

Collectively, these analyses demonstrate that REGAIN confidence provides a quantitative measure of heterogeneous predictability in histology-derived molecular inference. Although predictability varies across cell types, its variation across spatial domains is greater and persists within individual cell types, suggesting that local tissue context is more strongly associated with the reliability of molecular inference from H&E morphology than broad cellular identity.

### REGAIN confidence weighting enhances the prognostic value of histology-inferred spatial molecular features in breast cancer

To investigate the prognostic value of transcriptomic information inferred from histopathology, we performed survival analyses using the TCGA-BRCA cohort. Following cell segmentation quality control and clinical data matching, 766 patients with 119 overall survival events were included. Cell types were annotated from REGAIN-predicted single-cell gene expression profiles using SingleR, identifying nine major populations: B cells, myeloid cells, macrophages, stromal cells, malignant cells, T cells, plasma cells, endothelial cells, and epithelial cells. STAligner-based clustering of the predicted spatial transcriptomic profiles identified eight spatial domains (D0–D7), which were annotated according to their cell type composition and gene enrichment profiles as epithelial-enriched mixed (D0), immune–vascular stromal (D1), proliferative malignant (D2), proliferative and receptor-associated malignant (D3), immune-enriched epithelial (D4), receptor-associated malignant (D5), extracellular matrix–remodeling stromal (D6), and vascular-associated stromal (D7) domains.

Based on these annotations, we constructed six categories of spatial molecular features: (1) cell-type abundance, defined as the proportion of each cell type among all cells; (2) spatial-domain abundance, defined as the proportion of cells assigned to each spatial domain; (3) cell-type–domain joint abundance, defined as the proportion of all cells belonging to a specific cell type within a specific spatial domain; (4) cell-type distribution across spatial domains, defined as the proportion of cells of a given cell type assigned to each spatial domain; (5) cell-type composition within spatial domains, defined as the proportion of each cell type among all cells within a given spatial domain; and (6) spatial distribution diversity, quantified using entropy measures derived from the two complementary distributions defined in (4) and (5) (see Methods).

Each feature was evaluated in its original and confidence-weighted forms (see Methods), using age and PAM50 molecular subtype as the clinical baseline (Fig. 5a). Across most feature categories, confidence weighting generally yielded higher cross-validated Harrell’s concordance index (C-index) values and lower Cox P values compared with the corresponding unweighted features. Using Cox P < 0.05 and a mean cross-validated C-index exceeding the clinical baseline as screening criteria, 20 unweighted and 29 confidence-weighted features satisfied both thresholds. These findings suggest that confidence weighting strengthens the prognostic associations and predictive utility of histology-inferred spatial molecular features, thereby providing additional prognostic information beyond age and PAM50 molecular subtype.

**Fig. 5.**
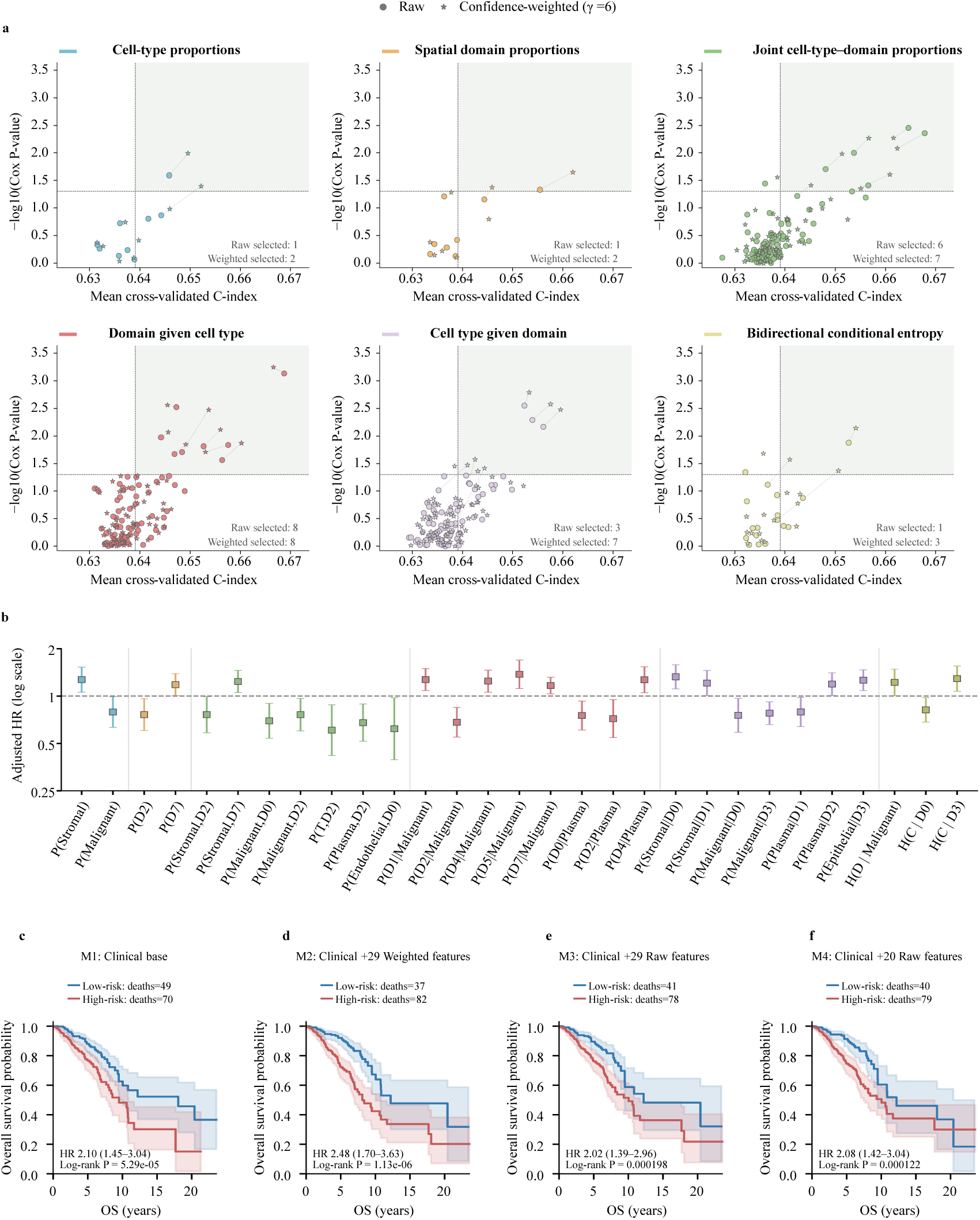
REGAIN confidence improves breast cancer survival risk stratification through spatial transcriptomic features. (a) Prognostic screening of 250 candidate cellular composition and spatial organization features across six categories, comparing original unweighted (circles) and confidence-weighted (stars) versions. The x-axis shows the mean cross-validated Harrell’s C-index, and the y-axis shows the P value from Cox regression adjusted for age and PAM50 molecular subtype. Short lines connect paired unweighted and confidence-weighted versions when at least one member of the pair meets both screening criteria (P < 0.05 and mean cross-validated C-index exceeding the clinical baseline); otherwise, the pair is not connected. (b) Forest plot showing hazard ratios (HRs) and 95% confidence intervals (CIs) for the 29 selected features, estimated using Cox regression adjusted for age and PAM50 subtype. The HR axis is displayed on a logarithmic scale. (c–f) Kaplan–Meier overall survival curves comparing high- and low-risk patient groups stratified using four Cox models: (c) age and PAM50 subtype alone, (d) age and PAM50 subtype combined with the 29 selected confidence-weighted features, (e) age and PAM50 subtype combined with the corresponding unweighted versions of the same 29 features, and (f) age and PAM50 subtype combined with 20 selected unweighted features.

We next examined the 29 selected confidence-weighted features in greater detail. The forest plot presents the individual features and their corresponding hazard ratios (HRs) and 95% confidence intervals (CIs), estimated from separate Cox regression models adjusted for age and PAM50 molecular subtype (Fig. 5b). Malignant-cell-related features were prominently represented among the selected features, consistent with the prognostic relevance of tumor cell abundance and spatial organization in breast cancer. In addition, features associated with plasma, stromal and epithelial cells highlighted the potential contributions of the tumor microenvironment to patient prognosis.

To evaluate patient risk stratification, we compared four Cox models: the clinical baseline comprising age and PAM50 subtype, the clinical baseline combined with the 29 selected confidence-weighted features, the clinical baseline combined with the unweighted versions of the same 29 features, and the clinical baseline combined with the 20 selected unweighted features. Kaplan–Meier analysis showed separation between high- and low-risk groups using the clinical baseline (HR = 2.10, 95% CI 1.45–3.04; log-rank P = 5.29 × 10⁻⁵; Fig. 5c). Incorporating the 29 confidence-weighted features resulted in greater separation between the risk groups (HR = 2.48, 95% CI 1.70–3.63; log-rank P = 1.13 × 10⁻⁶; Fig. 5d). In this model, 82 deaths occurred in the high-risk group compared with 37 in the low-risk group, whereas the corresponding numbers were 70 and 49 under the clinical baseline. By comparison, the model incorporating the unweighted versions of the same 29 features yielded weaker risk stratification than the clinical baseline (HR = 2.02, 95% CI 1.39–2.96; log-rank P = 1.98 × 10⁻⁴; Fig. 5e). Similarly, incorporating the 20 selected unweighted features did not improve risk stratification beyond the clinical baseline (HR = 2.08, 95% CI 1.42–3.04; log-rank P = 1.22 × 10⁻⁴; Fig. 5f).

Together, these findings demonstrate that prediction confidence contributes prognostically informative information beyond its role in assessing expression reconstruction reliability. By incorporating cell-level confidence into spatial feature construction, REGAIN identified a broader set of spatial molecular features associated with overall survival beyond age and PAM50 molecular subtype. Importantly, confidence-weighted features improved patient risk stratification beyond the clinical baseline, whereas their unweighted counterparts failed to provide comparable prognostic gains. These results establish prediction confidence as an important information dimension for extracting clinically informative spatial molecular patterns from routine histopathology, extending its value from assessing molecular predictability to capturing prognostically relevant tumor heterogeneity.

### REGAIN generalizes to spot-level spatial expression prediction

Although REGAIN was designed for single-cell-level spatial expression inference, its image-to-expression framework can also be adapted to spot-level spatial transcriptomics by replacing cell-level targets with spot-level expression profiles. We therefore evaluated REGAIN on the HER2^+^ breast cancer (HER2ST) spatial transcriptomics dataset^37^ using four-fold cross-validation, where H&E image patches were used to predict spot-level gene expression. Details of the four-fold data split are provided in the Methods.

REGAIN-predicted expression preserved spatially structured variation across representative HER2ST sections. In samples B1, B2, B3, G1, G2 and G3, the predicted expression supported spatial domain patterns that reflected visible histological heterogeneity in the corresponding H&E images (Fig. 6a, b). Similar spatial-domain patterns were also observed in additional HER2ST sections, including D1, D2 and D3 (Supplementary Fig. 11a), as well as H1, H2 and H3 (Supplementary Fig. 11b). These results suggest that REGAIN can capture morphology-associated spatial expression patterns beyond single-cell-resolution datasets.

**Fig. 6.**
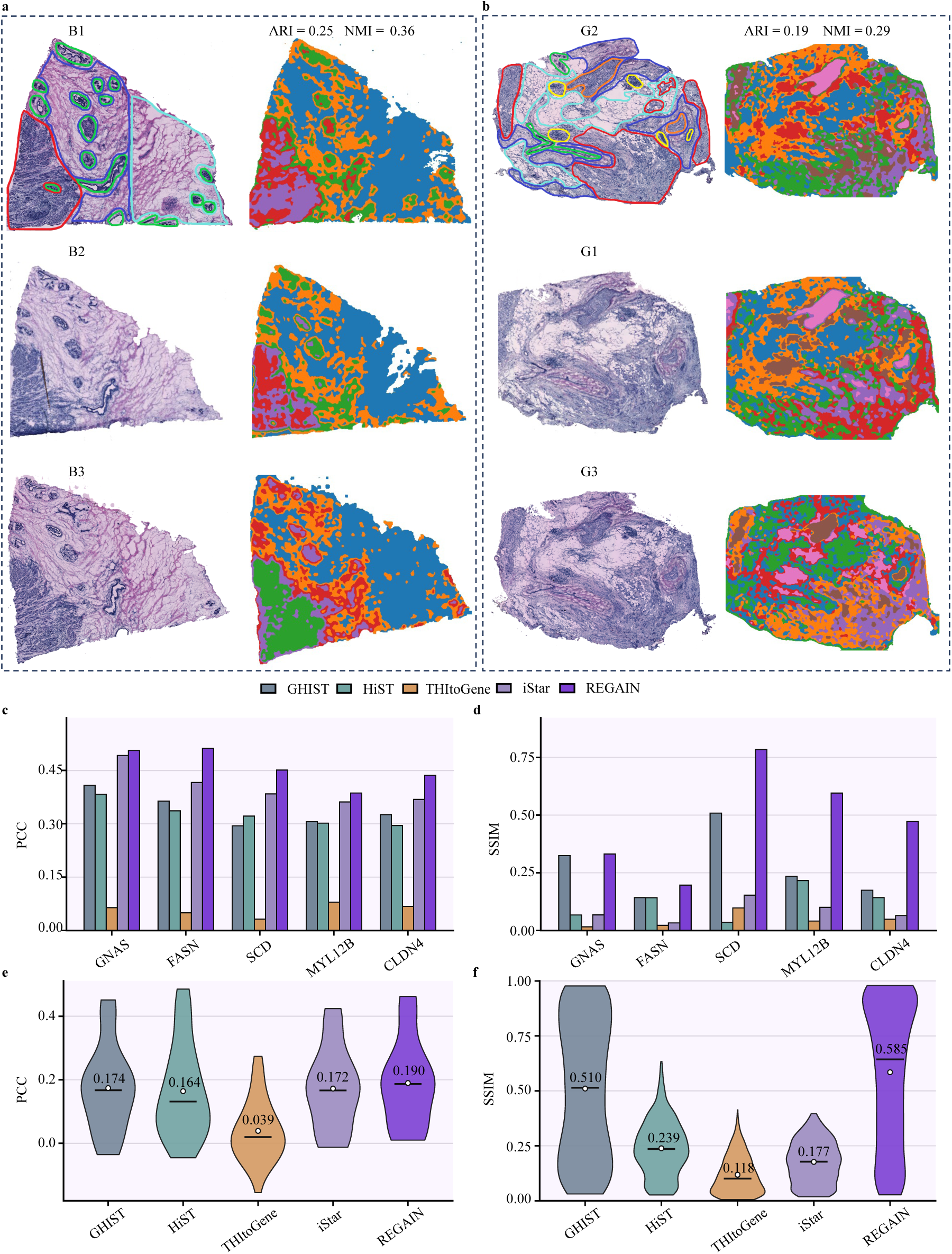
Spot-level spatial expression prediction on the HER2ST breast cancer dataset. (a,b) H&E images and spatial-domain identification results for representative HER2ST sections, including B1, B2, B3, G1, G2 and G3. For sections B1 and G2 with available spot-level spatial-domain annotations, ARI and NMI scores were calculated between the predicted domains and the ground-truth spatial domains after converting the inferred spatial domains to spot-level labels. (c) PCC comparison for marker genes across REGAIN and baseline H&E-to-ST prediction methods. (d) SSIM comparison for marker genes across methods. (e) PCC comparison across all genes. (f) SSIM comparison across all genes.

We next compared REGAIN with existing spot-level H&E-to-ST prediction methods, including iStar^11^, HiST^38^, THItoGene^39^ and GHIST^20^. REGAIN achieved the best performance for both marker genes and all genes in all metrics (Fig. 6c-f). Thus, although REGAIN was primarily designed for single-cell inference, its representation-learning framework remains effective in conventional spot-level spatial expression prediction.

Unlike the single-cell experiments, we did not perform confidence-guided filtering in this spot-level setting because the number of spots in each section is substantially smaller than the number of cells in single-cell datasets, making filtering less suitable for evaluation. Together, these results expand the applicability of REGAIN from single-cell spatial expression inference to spot-level H&E-to-ST prediction.

## Discussion

Histology-based spatial gene expression inference offers a promising approach to extending molecular characterization to large-scale pathology cohorts without requiring spatial transcriptomic profiling for every sample. However, its utility in precision medicine depends on two critical capabilities: generalizing morphology–expression relationships across tissue sections or datasets, and determining whether individual predictions are sufficiently reliable to support downstream biological and clinical analyses. REGAIN addresses these challenges by enabling single-cell spatial expression inference with improved cross-dataset generalizability and cell-level prediction confidence. Compared with spot-level prediction, single-cell resolution provides a more detailed representation of cellular heterogeneity, supporting cell-type annotation, spatial-domain identification and the characterization of tumor microenvironments. Importantly, by coupling expression prediction with reliability estimation, REGAIN provides a framework for extracting biologically meaningful spatial molecular information from routine H&E images, with potential applications in more accessible and scalable precision medicine.

To support generalizable single-cell expression inference, REGAIN integrates morphology– transcriptome contrastive pretraining with cell–patch joint supervision. Contrastive pretraining encourages the model to learn morphological representations associated with transcriptional states, while cell–patch joint supervision combines fine-grained cellular information with regional expression constraints, potentially reducing sensitivity to segmentation, image registration and cell-matching noise. Together, these strategies are designed to promote more robust morphology–expression representations across heterogeneous tissue samples. Consistent with this objective, REGAIN outperformed GHIST^20^ in cross-section and cross-dataset evaluations.

Beyond improving expression prediction, a central contribution of REGAIN is its ability to estimate the reliability of individual cell-level expression profiles. Our results demonstrate that high-confidence cells consistently exhibited more accurate expression recovery, while prediction reliability varied substantially across cells and tissue regions. This heterogeneity highlights the importance of incorporating confidence into downstream analyses. Confidence-guided analysis further improved the performance of both cell-type annotation and spatial-domain identification in cross-section and cross-dataset settings. Importantly, REGAIN confidence extends beyond conventional quality control by providing a reliability-aware framework for downstream biological interpretation. This extends the reliability problem raised by Stimage27 from uncertainty-aware histology-to-expression prediction to cell-level expression-profile reliability: rather than estimating uncertainty for individual gene predictions, REGAIN asks whether the entire predicted expression profile of each cell is reliable enough for downstream single-cell analysis.

We further investigated the biological significance of prediction confidence by examining variations in morphology–expression predictability across cell types and spatial domains in breast cancer tissues. In particular, malignant cells and malignant-epithelial regions generally exhibited higher confidence and more accurate expression recovery. Previous studies have shown that the transition from ductal carcinoma in situ to invasive breast cancer involves basement membrane breach, loss or disruption of the myoepithelial layer and stromal invasion^40^, accompanied by progressive changes in tumor–stroma structure and composition^41^. These histological changes may contribute to the distinctive morphological characteristics of malignant cells and strengthen their morphology–expression associations^42,43^, potentially explaining their higher prediction confidence and expression recovery accuracy. Importantly, prediction confidence also varied substantially across spatial domains even within the same cell type, suggesting that local tissue context influences morphology-associated predictability beyond cell-type identity. Such spatial heterogeneity may reflect differences in tissue architecture and microenvironmental composition, which can influence the morphological cues available for inferring gene expression.

The TCGA-BRCA survival analysis further suggests that REGAIN confidence may carry biologically and clinically relevant information beyond prediction reliability alone. As demonstrated by our analyses of cell types and spatial domains, confidence varies with morphology-associated transcriptional heterogeneity and spatial tissue organization, both of which are closely related to tumor progress. Notably, many of the confidence-weighted spatial features associated with survival beyond age and PAM50 molecular subtype were related to malignant cells, particularly their abundance and distribution across spatial domains. This finding highlights the prognostic relevance of tumor-associated spatial molecular heterogeneity captured through confidence-aware inference. Consistently, confidence weighting strengthened the prognostic associations of spatial molecular features and yielded more pronounced patient risk stratification than the corresponding unweighted features. Together, these findings suggest that prediction confidence may encode prognostically relevant biological information, rather than merely serving as a measure of expression prediction reliability.

Looking forward, the broader significance of REGAIN lies in introducing cell-level prediction reliability as an integral component of histology-based molecular inference. As computational methods increasingly enable molecular profiles to be inferred from routine H&E images, a fundamental challenge is no longer simply whether gene expression can be predicted, but whether these predictions can be trusted and meaningfully used for biological discovery and clinical applications. Rather than treating all inferred molecular profiles as equally reliable, future approaches could explicitly incorporate prediction confidence into the representation and analysis of histology-derived molecular data. Such a framework would enable prediction reliability to be propagated into downstream analyses, including cellular phenotyping, spatial organization, biomarker discovery and clinical risk modeling, while identifying tissue regions where direct molecular measurements remain essential. Importantly, this concept is not limited to REGAIN or transcriptomic prediction, but could be extended to other histology-based molecular inference tasks. By establishing prediction reliability as an intrinsic component of computationally inferred molecular information, REGAIN points toward a broader shift from accuracy-centered prediction to reliability-aware molecular characterization, potentially enabling more trustworthy and scalable precision medicine based on routine histopathology.

REGAIN still has limitations. Public single-cell-resolution H&E-spatial transcriptomics paired datasets remain limited, restricting the scale of validation. The model also depends on the quality of cell segmentation, image registration and cell matching. Although cell-patch joint supervision reduces sensitivity to local matching noise, segmentation and registration errors may still affect cell-level labels and downstream confidence estimation. In addition, the confidence score is an empirical reliability estimate, rather than a strictly calibrated probabilistic uncertainty. Cross-platform transfer in this study still required limited target-platform adaptation and did not represent zero-shot prediction. Future work should improve robustness to segmentation and registration errors and develop better calibrated uncertainty models. Overall, REGAIN provides a practical framework for reliability-aware single-cell molecular inference from routine histology, extending its utility from spatial molecular characterization to clinically relevant prognostic analysis.

## Acknowledgements

We thank all researchers and personnel involved in generating, curating and maintaining the public datasets used in the REGAIN study. Their efforts in data generation, annotation and preservation made this work possible.

This work was supported by the National Natural Science Foundation of China (No. 62572148) to Tianyi Zhao.

The funders had no role in study design, data collection and analysis, decision to publish or preparation of the manuscript.

## Author information

### Contributions

Xinyi Bai and Tianyi Zhao conceived the project and designed the overall study. Xinyi Bai performed the main experiments, developed the core analyses and drafted the manuscript framework. Renjie Liu conducted the multi-section spatial-domain analysis. Yuzhi Sun performed the spot-level comparative experiments. Tianyi Zhao provided conceptual guidance, critical suggestions and revised the manuscript. Guohua Wang and Yadong Wang jointly supervised the study, ensured the overall quality of the work and contributed to manuscript revisions. All authors have read and approved the final manuscript.

## Methods

### Datasets and preprocessing

#### Xenium datasets

We used three publicly available Xenium resources for single-cell spatial gene expression inference. The first breast cancer resource was the 10x Genomics human breast Xenium preview dataset, which contains two breast cancer samples and three tissue sections, including two consecutive sections from the first sample and one section from the second sample. The processed Xenium panels contained 313 genes for the first breast cancer sample and 280 genes for the second breast cancer sample. The second breast cancer resource was the 10x Genomics FFPE human breast dataset generated with the pre-designed Xenium Human Breast Gene Expression Panel, containing 280 genes. We also used a 10x Genomics FFPE human lung cancer Xenium dataset generated using a 377-gene Xenium Human Multi-Tissue and Cancer Panel, for which 10x Genomics provides cell segmentation outputs and cell-level expression matrices. These Xenium datasets provide post-Xenium H&E images, DAPI images, cell segmentation outputs and cell-level expression matrices.

For each Xenium section, the post-Xenium H&E image was aligned to the sequencing physical coordinate system using Xenium Explorer, following the 10x Genomics image-alignment workflow. The aligned H&E image was visually inspected to confirm the correspondence between histological structures and the Xenium/DAPI coordinate system. Cells were then segmented on the registered H&E image using Cellpose^44^. Because Xenium expression profiles are associated with cells defined in the DAPI/Xenium coordinate system, whereas REGAIN uses H&E-derived cellular morphology as input, we constructed paired H&E cell-expression samples by matching H&E-derived cell masks to Xenium/DAPI-derived cell masks.

For each H&E-derived cell, the Xenium/DAPI-derived cell with the largest spatial overlap was identified. A matched pair was retained when the overlapping area covered at least 50% of the H&E-derived cell mask area, following the label-construction strategy used in GHIST^20^. The expression profile and metadata of the matched Xenium/DAPI-derived cell were then assigned to the corresponding H&E-derived cell. Cells without a reliable match were excluded from model training and evaluation. This procedure produced paired H&E cell masks and single-cell spatial expression profiles for training REGAIN.

Gene expression profiles were normalized and log-transformed before model training. Registered H&E images were tiled into image patches, and all matched cells within each patch were collected together with their masks, coordinates and expression profiles. For cell-level supervision, the expression profile assigned to each H&E-derived cell was used as the single-cell target. For patch-level supervision, the expression profiles of all matched cells within the same patch were aggregated to generate patch-level expression targets.

#### Visium HD datasets

We used two publicly available 10x Genomics Visium HD datasets to evaluate Xenium-to-Visium HD transfer, including a human breast cancer Visium HD dataset and a human lung cancer Visium HD dataset. The breast cancer Visium HD dataset was paired with breast cancer Xenium data, and the lung cancer Visium HD dataset was paired with lung cancer Xenium data. These experiments were designed to assess whether morphology-transcriptome representations learned from Xenium could be adapted to Visium HD within the same cancer type.

For each Visium HD section, we extracted the 2 × 2 μm filtered barcode matrix, tissue-position parquet file and high-resolution H&E image from the Space Ranger output. H&E images were percentile-normalized before cell segmentation. Cells were segmented from the H&E image using Cellpose^44^, producing an instance-level mask image in which each cell was assigned a unique label. The mask image was then used directly for barcode-to-cell assignment.

Visium HD barcodes were mapped to the H&E image coordinate system according to the tissue-position file. A barcode was assigned to a cell if its spatial coordinate fell within a labeled cell region in the Cellpose^44^ mask image. Barcodes assigned to the same cell mask were aggregated to generate a cell-level expression profile. Cells were retained when the number of unique barcodes within the cell region was less than 2,000, the unique molecular identifier count was greater than 20 and the mitochondrial gene percentage was less than 15%. For model training, we used the gene sets processed following PRTS^19^, which retained differentially expressed genes associated with cell clusters and spatial cellular heterogeneity. This resulted in 1,656 genes for the lung cancer Visium HD dataset and 1,147 genes for the breast cancer Visium HD dataset. The resulting paired H&E-derived cell masks and Visium HD expression profiles were used for model adaptation and evaluation.

For cross-platform experiments, REGAIN was first trained on Xenium data and then fine-tuned using a small fraction of target-platform Visium HD data. The remaining Visium HD regions were held out for testing. Therefore, this experiment evaluated limited target-platform adaptation rather than zero-shot cross-platform prediction.

#### HER2ST spot-level dataset

To evaluate whether REGAIN could be extended to conventional spot-level spatial transcriptomics, we used the HER2ST dataset. This dataset contains 36 spatial transcriptomics sections from eight HER2^+^ breast cancer patients. Patients A-D each contributed six tissue sections, whereas patients E-H each contributed three tissue sections. Each section includes an H&E image, spatial spot coordinates and spot-level gene expression profiles.

HER2ST data were processed following the spot-level H&E-to-ST prediction setting used in GHIST^20^. For each sequencing spot, a 224 × 224 pixels H&E image patch centered on the spot was extracted and paired with the corresponding spot-level expression profile. Each patch was center-cropped to 112 × 112 pixels and used as the image input. For gene selection, the top 1,000 highly variable genes were first identified for each section, and genes expressed in fewer than 1,000 spots across all sections were removed, resulting in 785 genes for model training and evaluation.

In this experiment, REGAIN was adapted to the spot-level task by replacing cell-level expression targets with spot-level expression profiles. Four-fold cross-validation was used to evaluate model performance. To avoid information leakage across adjacent sections from the same patient, all sections from the same patient were assigned to the same fold. Thus, each fold contained sections from two patients, and the model was evaluated on held-out patients rather than randomly selected spots or sections. Confidence-guided filtering was not applied to the HER2ST experiment because each section contains far fewer spots than the number of cells available in single-cell Xenium datasets.

#### TCGA-BRCA dataset

Hematoxylin and eosin (H&E)-stained whole-slide images (WSIs) and corresponding clinical information were collected from The Cancer Genome Atlas Breast Invasive Carcinoma (TCGA-BRCA) cohort. The image download manifest was generated using the TCGAbiolinks R package, and WSIs were downloaded using the GDC Data Transfer Tool (gdc-client). Clinical information was retrieved using the TCGAretriever R package, while PAM50 molecular subtype annotations were obtained from the TCGA-BRCA phenotype data available through UCSC Xena. Tissue regions were extracted from the WSIs, and individual cells were segmented using Cellpose to obtain cell-level spatial coordinates and morphological regions. Following cell segmentation quality control and matching with clinical records, 766 patients with 119 overall survival events were retained for downstream clinical analysis.

### Overview of REGAIN

REGAIN was designed to infer single-cell spatial gene expression from H&E-stained histology images and to provide a cell-level reliability score for each prediction. Given an H&E image patch and the corresponding H&E-derived cell masks, REGAIN predicts the gene expression profile of each cell within the patch. In datasets with available cell type labels, REGAIN can additionally train a cell type annotation head, enabling direct cell type assignment during inference.

REGAIN adopts the main H&E-to-single-cell expression prediction framework of GHIST^20^ as the backbone, and introduces three key components to improve cross-modal representation learning, cell-level supervision and prediction reliability. First, morphology-transcriptome contrastive pretraining is used to align H&E patch representations with transcriptome-derived gene-symbol representations, thereby initializing a transcriptome-aware morphology encoder. Second, cell-patch joint supervision is introduced during single-cell expression prediction, combining cell-level expression loss with patch-level aggregate expression loss to reduce the effect of cell segmentation and H&E-DAPI matching errors. Third, a ridge-regression-based reliability estimator is trained to predict expression error from cell morphology embeddings and convert the predicted error into a confidence score. Through these components, REGAIN outputs both single-cell gene expression predictions and cell-level confidence estimates from H&E images.

#### Morphology-transcriptome contrastive pretraining

To initialize the histology encoder with transcriptome-aware representations, we performed morphology-transcriptome contrastive pretraining before training the single-cell expression prediction model. Each training instance consisted of an H&E image patch and the expression profiles of matched cells located within the patch. For the *i*-th patch, let **X***_i_* ∈ ℝ*^ni^*^×*G*^ denote the expression matrix of the *n_i_* matched cells within the patch, where *G* is the number of genes. We first aggregated cell-level expression profiles within the patch to obtain a patch-level expression vector,

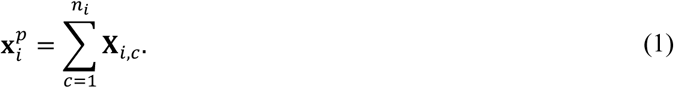

The 50 genes with the highest values in *x^p^_i_* were selected, and their gene symbols were concatenated into an expression-derived gene-symbol sentence *s_i_* . This sentence was used as a compact transcriptomic description of the dominant expression state of the patch. The H&E patch was encoded using an image encoder following the main histology feature extraction backbone of GHIST^20^. The same encoder architecture was subsequently used as the H&E feature extractor in the single-cell expression prediction model. Given an H&E patch **I***_i_*, the image backbone extracted multi-scale histology feature maps,

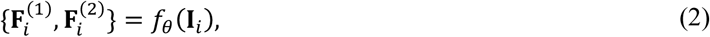

where **F***_i_*^(1)^and **F***_i_*^(2)^ denote feature maps from two resolution levels of the backbone. Patch-level features were obtained by global average pooling over each feature map,

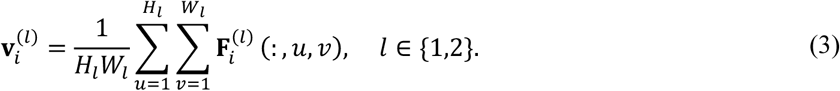

The pooled multi-scale features were concatenated and passed through a trainable image projection head *g_φI_* to obtain the patch-level morphology embedding,

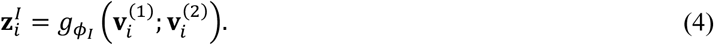

For the transcriptome branch, the gene-symbol sentence *s_i_* was encoded using a CoCa^23^ pretrained text encoder implemented with OpenCLIP^45^. Let *t_φ_* denote this text encoder. The sentence embedding was computed as

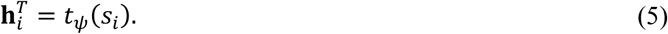

The text embedding was then mapped to the shared contrastive embedding space using a trainable two-layer projection head *q_ωT_*,

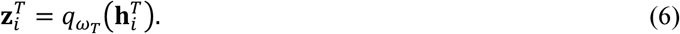

Both image and text embeddings were L2-normalized before contrastive learning,

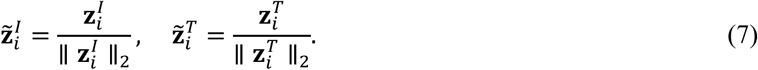

For a mini-batch of *B* matched H&E patches and gene-symbol sentences, the image-text similarity matrix was computed as

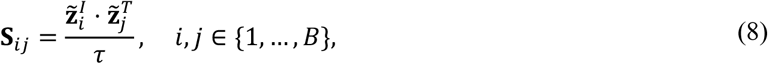

where *τ* is the temperature parameter. Matched H&E patches and gene-symbol sentences with the same batch index were treated as positive pairs, whereas all non-matched pairs in the same mini-batch were treated as negatives. We optimized a bidirectional contrastive loss consisting of image-to-text and text-to-image cross-entropy terms,

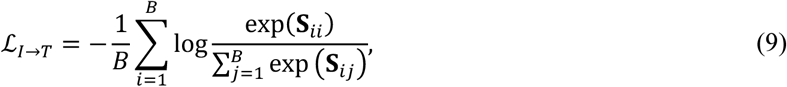

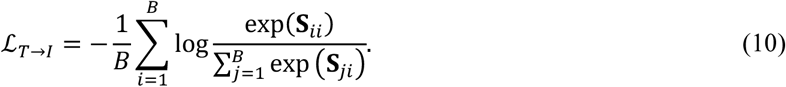

### The final contrastive pretraining objective was

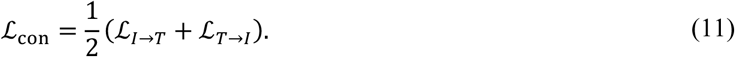

After contrastive pretraining, the learned image encoder parameters were used to initialize the H&E feature extractor in the downstream single-cell expression prediction model.

#### Single-cell expression prediction with cell-patch joint supervision

After morphology-transcriptome contrastive pretraining, REGAIN was trained to predict single-cell gene expression from H&E image patches. The expression prediction model used the main image-to-single-cell prediction framework of GHIST^20^, with the H&E feature extractor initialized by the contrastively pretrained image encoder. Given an H&E patch **I***_b_*, the image encoder extracted multi-scale histology feature maps,

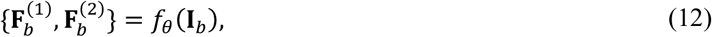

where 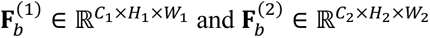 denote two feature maps from different resolution levels. For the *i*-th cell in patch *b*, its H&E-derived cell mask was resized to the spatial resolution of each feature map, resulting in 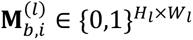 . Cell-level local morphology features were obtained by mask-based average pooling,

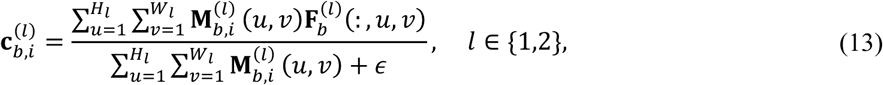

where *ε* is a small constant for numerical stability. In parallel, patch-level global morphology features were obtained by average pooling over the entire feature maps,

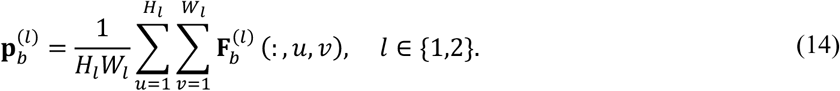

For each cell, the local cell features and global patch features were concatenated and mapped to a cell morphology embedding,

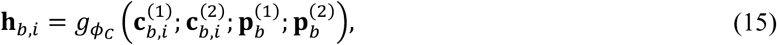

where *g_φC_* denotes the histology embedding module. The expression prediction head *r_ωE_* then mapped the cell morphology embedding to a gene expression vector,

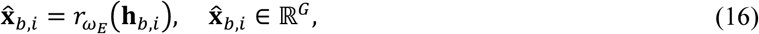

where *G* is the number of predicted genes. The cell-level expression loss was defined as the mean squared error between the predicted expression profile and the matched single-cell expression label,

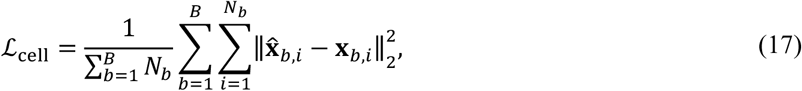

where *B* is the batch size, *N_b_* is the number of matched cells in patch *b*, and **x***_b_*_,*i*_ denotes the measured expression profile assigned to the *i*-th H&E-derived cell.

To reduce the influence of cell segmentation errors, H&E-DAPI registration errors and imperfect cell matching, REGAIN further introduced a patch-level aggregate expression constraint. For each patch, the patch-level expression target was computed by summing the expression profiles of all matched cells within that patch,

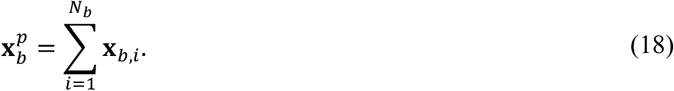

In parallel to the cell-level expression prediction branch, a patch-level prediction branch used the global patch morphology features to predict the aggregate expression profile of the patch. Specifically, the multi-scale global features were mapped to a patch morphology embedding,

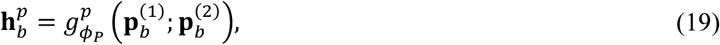

and the patch-level expression prediction head *r^p^* produced the aggregate expression prediction,

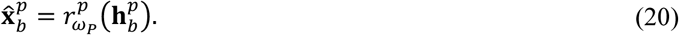

The patch-level loss was defined as the mean squared error between the predicted patch-level expression and the aggregated expression target,

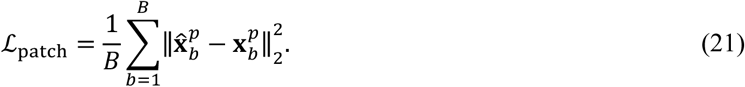

When cell type annotations were available, a cell type annotation head was trained together with the expression prediction model. The classifier *a_η_* produced cell type logits from the morphology embedding,

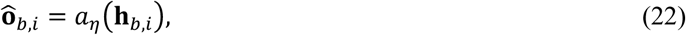

and the corresponding classification loss was denoted as ℒ_ct_.

The final training objective was

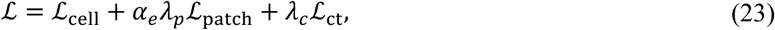

where *λ_p_* and *λ_c_* are loss weights. The coefficient *α_e_* indicates whether the patch-level loss was applied at training epoch *e*. In our implementation, the patch-level loss was applied during the first two epochs and then every two epochs thereafter. This intermittent strategy provided regional expression supervision while maintaining cell-level expression prediction as the primary optimization objective.

#### Reliability estimation and confidence-guided inference

REGAIN estimates a reliability score for each cell-level expression prediction. The reliability estimator was trained to predict the expected expression error from the H&E-derived cell morphology embedding. To construct the training set for reliability estimation, we held out a subset of cells from the training data. The single-cell expression prediction model was trained on the remaining cells and then applied to the held-out cells to obtain both the predicted expression profiles and the corresponding morphology embeddings.

For the *i*-th held-out cell, let **h***_i_* ∈ ℝ*^D^* denote the morphology embedding produced by the H&E encoder and histology embedding module, and let **x**a*_i_* ∈ ℝ*^G^* and **x***_i_* ∈ ℝ*^G^* denote the predicted and measured expression profiles, respectively. The normalized absolute error of gene *g* in cell *i* was computed as

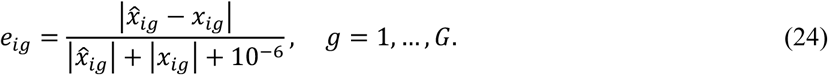

The gene-wise errors were averaged across all predicted genes to obtain a scalar cell-level expression error,

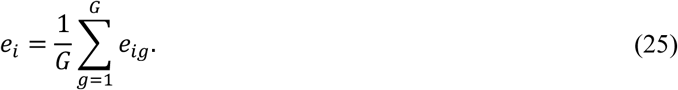

This produced a reliability training set 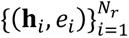 , where *N_r_* denotes the number of held-out cells used for reliability estimation.

Before training the reliability model, the cell morphology embeddings were standardized feature-wise using the mean and standard deviation estimated from the reliability training set,

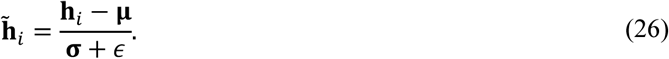

We then trained a ridge regression^46^ model to predict the scalar expression error from the standardized morphology embedding. The ridge model is defined as

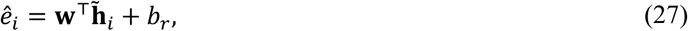

where **w** and *b*ᵣ are the regression coefficients and ridge-regression intercept. The model parameters were estimated by minimizing the squared prediction error with an *L*_2_ regularization term,

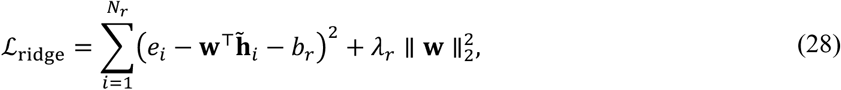

where *λ_r_* is the ridge regularization coefficient. During inference, REGAIN first predicted the single-cell expression profile and morphology embedding for each H&E-derived cell. The morphology embedding was standardized using the same scaler fitted on the reliability training set and then passed to the trained ridge regression model to estimate the expected cell-level expression error,

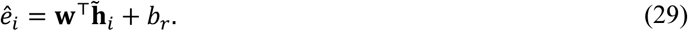

The confidence score was defined as an exponential transformation of the predicted error,

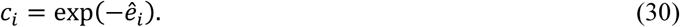

Thus, cells with lower predicted expression errors receive higher confidence scores. The confidence score was used to rank cell-level predictions, perform confidence-guided filtering and assess spatial variation in prediction reliability. This score represents an estimated reliability of the predicted expression profile rather than a calibrated probabilistic uncertainty.

### Cell type annotation and spatial domain analysis

#### Cell type annotation

Cell type annotation is a critical step in single-cell transcriptomic analysis. Existing approaches mainly rely on marker-gene-based annotation or reference transcriptome-based classification strategies, including scType^47^, SingleR^48^, scClassify^49^ and clustifyr^50^. In REGAIN, except for the TCGA-BRCA dataset, cell type annotation was performed based on the predicted single-cell expression profiles using a supervised neural-network-based classifier, following the strategy of recent histology-to-transcriptome frameworks such as SpatialEx^21^. Specifically, REGAIN used a cell type prediction head trained together with the single-cell expression prediction model. The annotation head took the learned cell morphology embedding as input. For the *i*-th cell in patch *b*, the morphology embedding **h***_b_*_,*i*_ was mapped to cell type logits by a classifier *a_η_*,

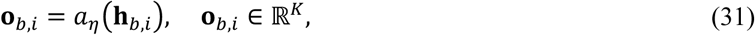

where *K* denotes the number of cell type categories. The predicted cell type probability was computed using a softmax function,

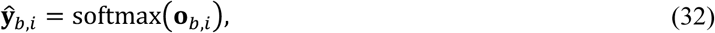

and the predicted cell type was assigned as

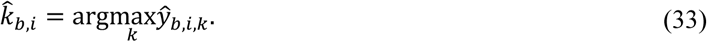

When cell type labels were available for the source training data, the annotation head was optimized using a cross-entropy loss,

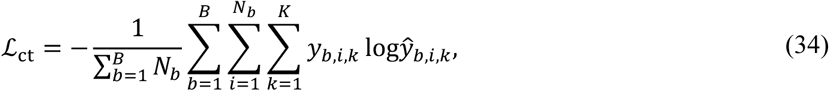

where *y_b_*_,*i*,*k*_ denotes the one-hot cell type label. In the cross-section experiment, the annotation head was trained on the annotated source Xenium breast section and applied to the target section together with expression prediction. In the cross-dataset experiment, the annotation head was trained on the annotated source Xenium breast dataset and then applied to the target dataset. For target datasets without public cell type annotations, reference labels for evaluation were generated from the measured expression profiles using the same expression-based annotation procedure. Thus, cell type prediction was evaluated against either public annotations or expression-derived reference labels, depending on the availability of manual or public cell type labels.

#### Spatial domain analysis

For the BreastCancer1 Rep2 and TENX94 datasets, spatial domains were identified from measured or predicted cell-level expression profiles using STAligner^36^. STAligner^36^ was first trained to obtain low-dimensional cell embeddings, denoted by *z^SD^* . The embeddings were aggregated onto an H&E-aligned spatial grid and spatially smoothed within the tissue foreground. The normalized grid embeddings and spatial coordinates were then combined for MiniBatchKMeans clustering^51^:

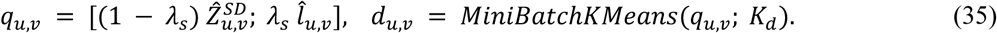

Here, 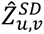 is the normalized, spatially smoothed embedding at grid location (u, v), *l_uv_* is its normalized spatial coordinate, and *λ_s_* = 0.1 is the location weight. We used *K_d_* = 8 spatial domains and a fixed random seed. The superscript SD denotes spatial domains. Grid-level domain labels were mapped back to individual cells, and domains inferred from predicted expression were compared with those derived from measured expression using the ARI and NMI.

In the confidence-guided pipeline, cell-level prediction confidence *c_i_* ∈ [0, 1] was converted to a reliability weight *ρ_i_* = (1 − *λ_a_*) + *λ_a_ c_i_*, where *λ_a_* = 0.5 controls the contribution of confidence. These weights modulated STAligner graph attention during training and embedding extraction:

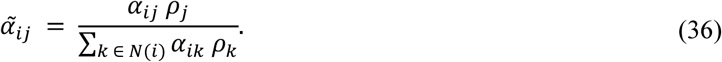

Here, *α_i_*_j_ denotes the attention weight from neighboring cell j to target cell i before confidence weighting, and N(i) denotes the graph neighborhood of cell i. More reliable neighbors therefore contributed more strongly to the learned embedding. Cell embeddings within each grid were subsequently aggregated using the same reliability weights:

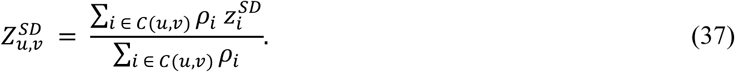

Here, C(u, v) is the set of cells assigned to grid location (u, v). During subsequent Gaussian spatial smoothing, contributions from observed grids were weighted by reliability calculated from their mean cell confidence. Interpolated grids were not treated as additional observed sources. The resulting grid representations were combined with spatial coordinates and clustered as described above. Small isolated regions were subsequently merged to improve spatial continuity.

We further refined the initial spatial domain assignments using the spatial distribution of prediction confidence. Cell-level confidence scores were mapped to the spatial grid and smoothed to form a local confidence map, from which large connected regions of elevated confidence were extracted. Within each region, the dominant domain was identified as the label occupying the largest grid area in the original clustering result, and all grids in that region were reassigned to this label. Labels outside these regions or at locations with insufficient confidence support were retained.

For the HER2ST spot-level spatial transcriptomics dataset, due to the sparsity of spot-level measurements compared with single-cell datasets, we directly applied clustering to the spatial feature representations extracted before expression prediction. The resulting spatial domains were mapped to the corresponding spots and compared with available spot-level domain annotations using ARI and NMI.

For the remaining spatial domain analyses, our primary aim was to obtain cell-level spatial domain labels for downstream analyses. We therefore used the standard STAligner^36^ spatial domain identification workflow.

### Clinical survival analysis

To evaluate the clinical relevance of REGAIN-inferred spatial transcriptomic profiles, we applied the REGAIN model trained on the BreastCancer2 Xenium dataset to H&E images from the TCGA-BRCA cohort to predict single-cell gene expression profiles and corresponding confidence scores. Cell types were annotated using SingleR with the breast cancer single-cell RNA-seq reference dataset (GSE17607852). Spatial domains were identified using STAligner36 based on predicted expression profiles and cell spatial coordinates. Eight spatial domains (D0–D7) were identified and characterized according to their cell type composition and gene enrichment patterns.

Construction of cellular composition and spatial organization features. For each patient, only successfully matched cells with known cell type annotations were retained; cells labeled Unknown were excluded. Let *N* denote the number of retained cells, *C_i_* ∈ {1, … ,9} the cell type of cell *i*, and *D_i_* ∈ {0, … ,7} its spatial domain. Let *N_c_*, *N_d_* and *N_c_*_,*d*_ denote the numbers of retained cells belonging to type *c*, domain *d* and their intersection, respectively. We constructed 250 candidate features across six categories.

Cell type proportions (9 features):

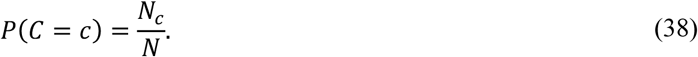

Spatial domain proportions (8 features):

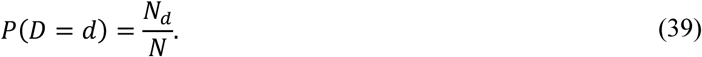

Joint cell type–domain proportions (72 features):

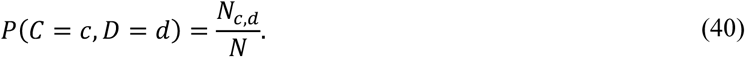

Domain distributions conditional on cell type (72 features):

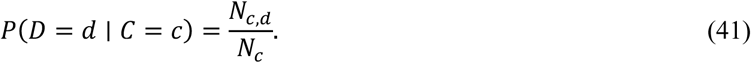

Cell type compositions conditional on domain (72 features):

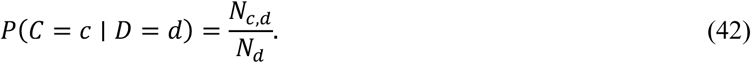

Normalized conditional entropies of domain assignments for each cell type and cell type compositions within each domain (17 features: 9 cell type-wise and 8 domain-wise):

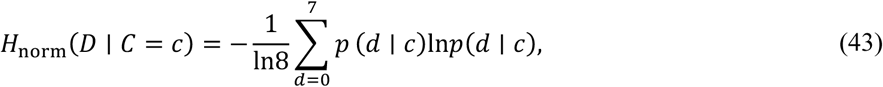

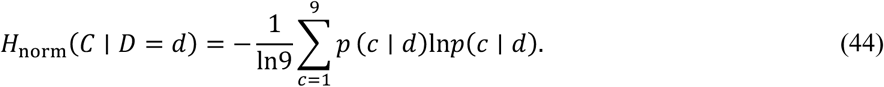

Here, *p*(*d* ∣ *c*) = *P*(*D* = *d* ∣ *C* = *c*); natural logarithms were used with the convention 0ln0 = 0 . The entropy ranges from 0 (all cells of the type occupy one domain) to 1 (uniform distribution across eight domains). The normalization denominator remained ln8 even when fewer than eight domains were present in a patient. For the eight domain-wise entropy features, *p*(*c* ∣ *d*) = *P*(*C* = *c* ∣ *D* = *d*). Values range from 0 (all cells in the domain belong to one cell type) to 1 (uniform composition across nine cell types), with a fixed normalization denominator of ln9. Entropies with zero conditioning denominators were treated as missing.

Confidence-weighted features. Each candidate feature was also calculated in a confidence-weighted form. Let *q_i_* denote the raw REGAIN confidence score for cell *i* and *q*_max_ the maximum confidence among matched, retained cells from the same patient. We used the patient-wise maximum-normalized sixth-power weight:

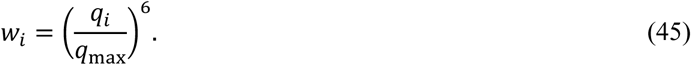

For example, the confidence-weighted joint cell type–domain proportion was defined as

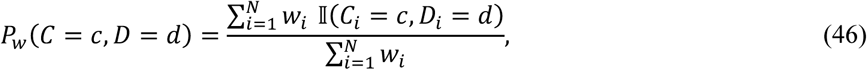

and the confidence-weighted domain distribution conditional on cell type as

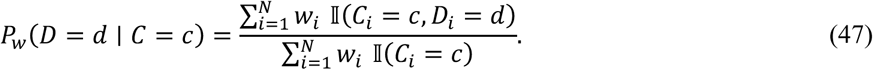

Here I(⋅) denotes an indicator function. All other weighted proportions used the corresponding weighted counts and denominators. Both weighted entropy measures were calculated from the corresponding weighted conditional probabilities.

Survival modeling and evaluation. Prognostic associations were assessed using Cox proportional hazards regression adjusted for age and PAM50 molecular subtype, alongside cross-validated Harrell’s concordance index (C-index). Candidate features were screened using a Cox regression P value < 0.05 and a mean cross-validated C-index exceeding that of the clinical baseline, yielding 29 confidence-weighted and 20 unweighted features. Four Cox models were compared: age and PAM50 subtype alone, the clinical baseline combined with the 29 selected confidence-weighted features, the clinical baseline combined with the corresponding unweighted versions of the same 29 features, and the clinical baseline combined with the 20 independently selected unweighted features. Kaplan–Meier curves and log-rank tests were used to compare overall survival between high- and low-risk patient groups.

### Evaluation metrics

We evaluated REGAIN and baseline methods using expression prediction metrics, spatial pattern similarity, downstream annotation performance, spatial domain consistency and morphology-associated separability.

For gene-level expression prediction, we computed the Pearson correlation coefficient (PCC) between predicted and measured expression values across cells or spots. For gene *g*, PCC was defined as

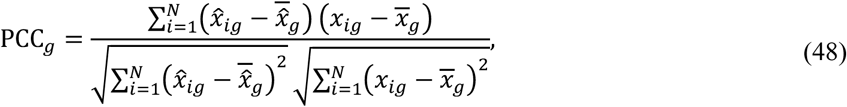

where *x_i_*_g_ and *x*a*_i_*_g_ denote the measured and predicted expression values of gene *g* in cell or spot *i* , respectively, and *N* is the number of evaluated cells or spots. Gene-wise PCC values were averaged across genes to summarize overall prediction performance.

We further computed the root mean squared error (RMSE) to quantify the absolute prediction error between predicted and measured expression profiles,

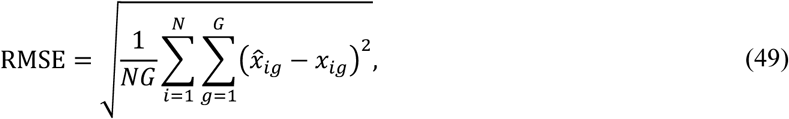

where *G* denotes the number of evaluated genes.

To evaluate whether spatial expression patterns were preserved, we computed the structural similarity index measure (SSIM) between predicted and measured spatial expression maps. For each gene, expression values were projected to their spatial coordinates to generate paired predicted and measured expression maps. SSIM was computed as

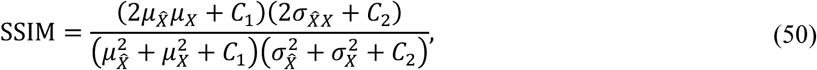

where (X) and *X*^{^ denote the measured and predicted expression maps, *μ* , *σ*^2^ and *σ_X̂_*_*X*_ denote the corresponding means, variances and covariance, and *C*_1_ and *C*_2_ are constants for numerical stability.

In addition to gene-wise metrics, we computed cell-wise PCC to evaluate whether the relative expression pattern across genes was preserved within each cell. For cell *i*, cell-wise PCC was defined as

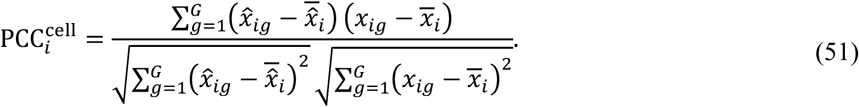

The final cell-wise PCC was reported as the average or median of PCC^cell^ across evaluated cells, depending on the analysis.

Cell type annotation performance was evaluated using annotation accuracy. Given the predicted cell type label *y*a*_i_* and the reference label *y_i_*, accuracy was computed as

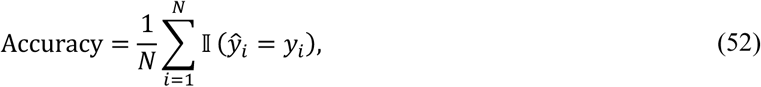

where I(⋅) is the indicator function. Reference labels were obtained from public cell type annotations when available, or from expression-derived annotation labels generated using measured expression profiles.

Spatial domain consistency was evaluated using ARI and NMI. ARI measures the agreement between predicted and reference spatial domain assignments after correcting for chance, whereas NMI quantifies the normalized mutual dependence between the two cluster labelings. Both metrics were computed between spatial domains inferred from predicted expression profiles and reference domains inferred from measured expression profiles.

To quantify morphology-associated separability of different cell groups, we computed a separability score in the standardized morphology embedding space. Let **h***_i_* denote the standardized morphology embedding of cell *i*, and let G*_k_* denote the set of cells belonging to group *k*. The centroid of group *k* was computed as

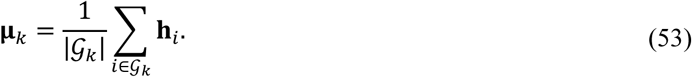

The intra-group distance was defined as the average Euclidean distance between cells in group *k* and their group centroid,

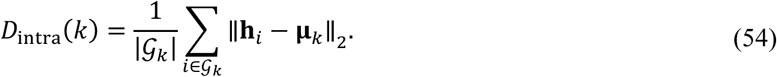

The inter-group distance was defined as the average Euclidean distance between the centroid of group *k* and the centroids of all other groups,

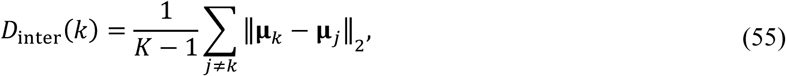

where *K* is the number of cell groups. The separability score of group *k* was then computed as

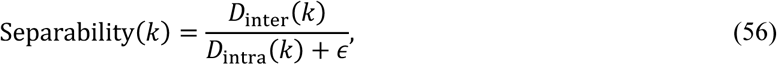

where *ε* is a small constant for numerical stability. A higher separability score indicates that a cell group is more distant from other groups while being more compact within its own group in the morphology embedding space. We used this metric to analyze the relationship between morphology-derived group distinctiveness, prediction confidence and cell-wise prediction accuracy.

### Implementation details

REGAIN was implemented in Python 3.10.18 using PyTorch 2.1.1 with CUDA 11.8. All experiments were performed on NVIDIA L20 GPUs with 46 GB memory. Unless otherwise specified, H&E images were divided into 256 × 256 patches, and each patch contained at most 200 cells. Stain augmentation was applied during training to improve robustness to H&E staining variation.

The expression prediction model used a morphology embedding dimension of 256 and a dropout rate of 0.05. The model was trained for 50 epochs with a batch size of 8. We used the Adam optimizer with an initial learning rate of 1 × 10^−3^, *β*_1_ = 0.9, *β*_2_ = 0.999, *ε* = 1 × 10^−8^ and weight decay of 1 × 10^−4^. The model checkpoint was saved after each epoch. Cell type annotation was enabled when cell type labels were available in the training data.

For the single-cell expression prediction model, the H&E feature extractor was initialized using the image encoder obtained from morphology-transcriptome contrastive pretraining. The downstream model was trained with the combined objective of cell-level expression prediction, patch-level aggregate expression prediction and, when applicable, cell type annotation. During inference, the trained model generated cell-level expression profiles and morphology embeddings for H&E-derived cells. These embeddings were further used by the ridge-regression reliability estimator to compute cell-level confidence scores.

## Data availability

All datasets used in this study are publicly available. The breast cancer Xenium preview dataset is available from 10x Genomics at https://www.10xgenomics.com/products/xenium-in-situ/preview-dataset-human-breast. The FFPE human breast Xenium dataset is available at https://www.10xgenomics.com/datasets/ffpe-human-breast-with-pre-designed-panel-1-standard. The FFPE human lung cancer Xenium dataset is available at https://www.10xgenomics.com/datasets/preview-data-ffpe-human-lung-cancer-with-xenium-multimodal-cell-segmentation-1-standard. The Visium HD human breast cancer dataset is available at https://www.10xgenomics.com/datasets/visium-hd-cytassist-gene-expression-human-breast-cancer-fixed-frozen, and the Visium HD human lung cancer dataset is available at https://www.10xgenomics.com/datasets/visium-hd-cytassist-gene-expression-human-lung-cancer-post-xenium-expt. The HER2ST dataset is available at https://github.com/almaan/her2st. The TCGA-BRCA H&E whole-slide images are available through the Genomic Data Commons (GDC) Data Portal at https://portal.gdc.cancer.gov/projects/TCGA-BRCA. TCGA-BRCA clinical data are available through cBioPortal at https://www.cbioportal.org/study/clinicalData?id=brca_tcga_pub, and PAM50 molecular subtype annotations were obtained from the TCGA-BRCA phenotype data available through UCSC Xena at https://xena.ucsc.edu/.

## Code availability

The source code of REGAIN is available at https://github.com/Bxinyi2025/REGAIN, including the implementations of morphology-transcriptome contrastive pretraining, single-cell spatial gene expression prediction and confidence estimation.

