## Supplementary notes and figures for "Confidence-guided spatial gene expression inference for reliability-aware prognostic analysis from histology"

### for

**Supplementary Note 1.** REGAIN enables cross-platform expression inference transfer across cancer types.

**Supplementary Fig 1:** Additional marker-gene visualization for the rep2 cross-section test sample.

**Supplementary Fig 2:** Cell-type composition of the Xenium breast rep2 cross-section test sample after confidence-guided filtering.

**Supplementary Fig 3:** Spatial distribution of cell-level prediction confidence in the breast1rep2 cross-section test sample.

**Supplementary Fig 4:** Comparison of ground-truth and REGAIN-predicted cell-type distributions in the cross-section breast cancer dataset.

**Supplementary Fig 5:** Spatial visualization of representative marker genes in the TENX94 cross-dataset test sample.

**Supplementary Fig. 6:** Spatial visualization of additional breast cancer-associated marker genes in the TENX94 cross-dataset test sample.

**Supplementary Fig 7:** Cell-type composition changes under confidence-guided filtering in the TENX94 cross-dataset test sample.

**Supplementary Fig 8:** Spatial distribution of cell-level prediction confidence in the TENX94 cross-dataset test sample.

**Supplementary Fig 9:** Xenium-to-Visium HD cross-platform transfer in breast and lung cancers.

**Supplementary Fig 10:** Additional spatial-domain results on the HER2ST spot-level dataset.

**Supplementary Note 1. REGAIN enables cross-platform expression inference transfer across cancer types**

We next evaluated whether REGAIN could transfer morphology-expression knowledge across spatial transcriptomic platforms. We focused on Xenium-to-Visium HD transfer as a more challenging cross-platform generalization setting, because these platforms differ substantially in spatial resolution, capture chemistry, gene coverage and measurement characteristics. We performed this evaluation separately in breast cancer and lung cancer. In each cancer type, REGAIN was first trained on Xenium data and then adapted using a small fraction of Visium HD data before prediction on held-out Visium HD regions, thereby evaluating limited target-platform adaptation rather than zero-shot transfer.

In the breast cancer experiment, the Xenium source data were derived from BreastCancer1 Rep1. The target Visium HD breast cancer dataset contained 285,594 matched cells after preprocessing, and the processed gene panel contained 1,147 genes. We used 20% of the Visium HD regions for target-platform fine-tuning, resulting in 22,985 training cells. Among these cells, 20% were further held out to train the confidence model. The remaining 262,609 matched cells were used for final prediction and evaluation. In the lung cancer experiment, the Xenium source data were derived from the 10x Genomics FFPE human lung cancer Xenium dataset, which contained 106,146 matched cells and a 377-gene panel. The target Visium HD lung cancer dataset contained 94,085 matched cells after preprocessing, and the processed gene panel contained 1,656 genes. Similarly, 20% of the Visium HD section was used for fine-tuning, resulting in 20,330 training cells, of which 20% were further used to train the confidence model. The remaining 73,755 matched cells were used for final prediction and evaluation.

In breast cancer, REGAIN outperformed GHIST in recovering Visium HD-derived expression from H&E images across all evaluated gene sets, including all genes, HVGs and SVGs (Supplementary Fig. 9a). Confidence-guided filtering further improved prediction quality, increasing gene-wise SSIM and reducing cell-wise RMSE among high-confidence spatial units (Supplementary Fig. 9b, c). Spatial visualization of representative breast cancer marker genes, including PGR and EPCAM, showed that REGAIN preserved major expression patterns in held-out Visium HD regions (Supplementary Fig. 9d).

We then tested whether the same transfer strategy generalized to another cancer type. In lung cancer, REGAIN achieved higher PCC than GHIST across all gene sets in the Xenium-to-Visium HD transfer setting (Supplementary Fig. 9e). Confidence-guided selection again enriched higher-quality predictions, as reflected by improved gene-wise SSIM and reduced RMSE after retaining high-confidence spatial units (Supplementary Fig. 9f, g). Spatial visualization of representative lung cancer marker genes, including KRT8 and EPCAM, further showed that REGAIN preserved spatial expression structure in held-out Visium HD regions (Supplementary Fig. 9h).

Together, these results show that REGAIN can transfer histology-to-expression inference across cancer types. Although this experiment does not represent zero-shot cross-platform prediction, it demonstrates that REGAIN provides an effective warm-start framework for transferring morphology-transcriptome representations across spatial transcriptomic platforms with limited target-platform supervision.

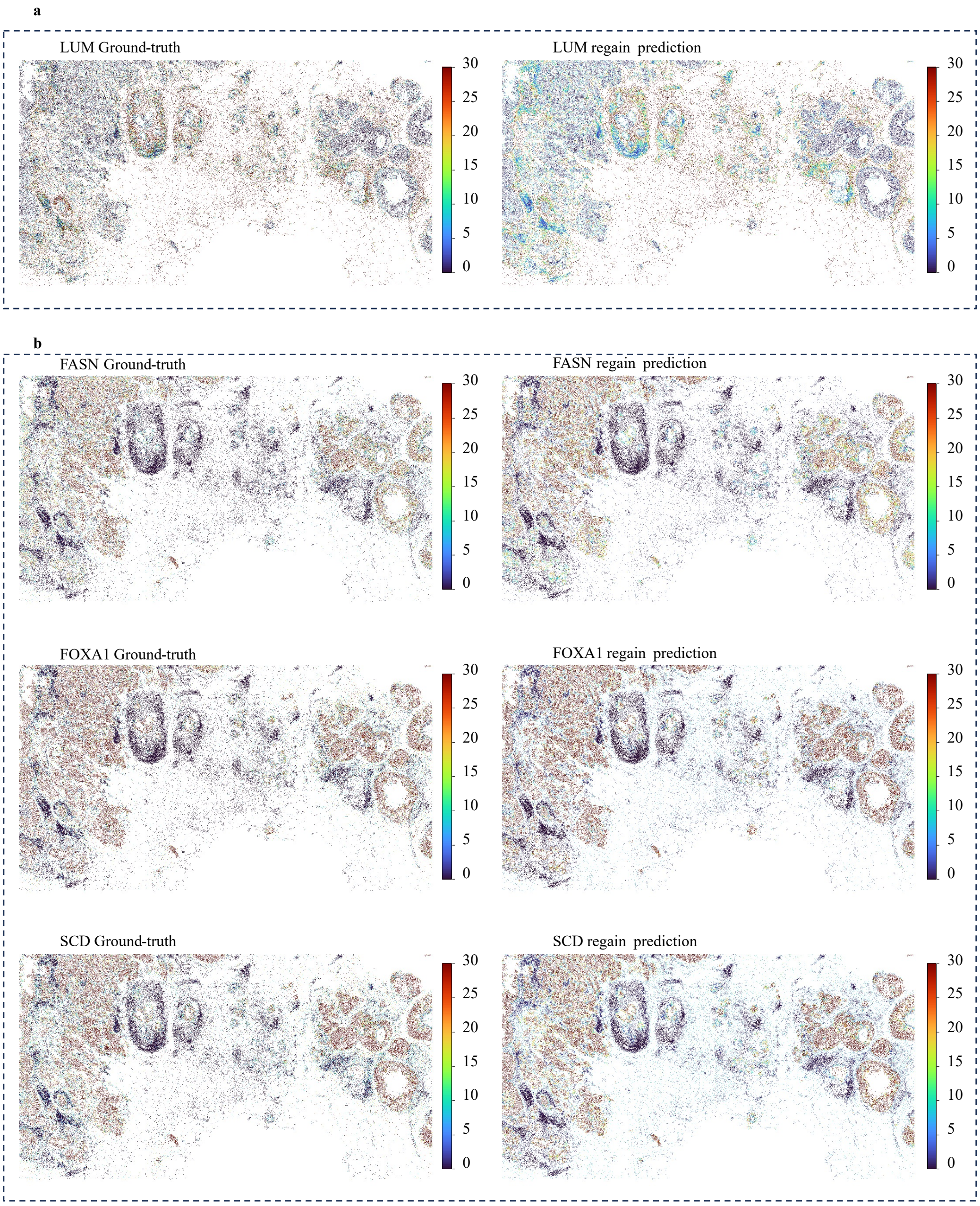


**Supplementary Fig. 1 | Additional marker-gene visualization for the rep2 cross-section test sample.** (a) Spatial expression patterns of the stromal marker LUM. (b) Spatial expression patterns of additional breast cancer- or tumor-associated marker genes.

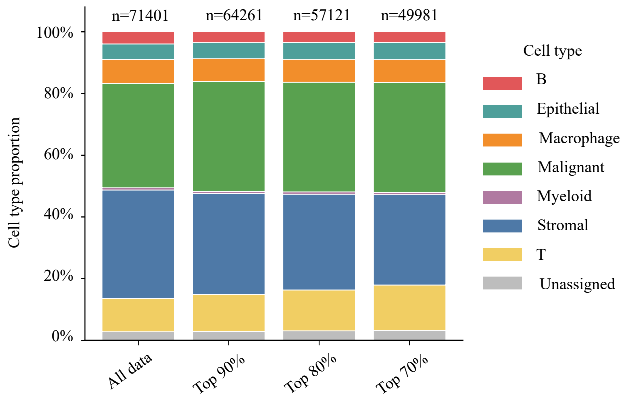


**Supplementary Fig. 2 | Cell-type composition of the Xenium breast rep2 cross-section test sample after confidence-guided filtering.** The x-axis shows all cells and subsets retained after confidence-based selection, enabling comparison of cell-type proportions before and after filtering.

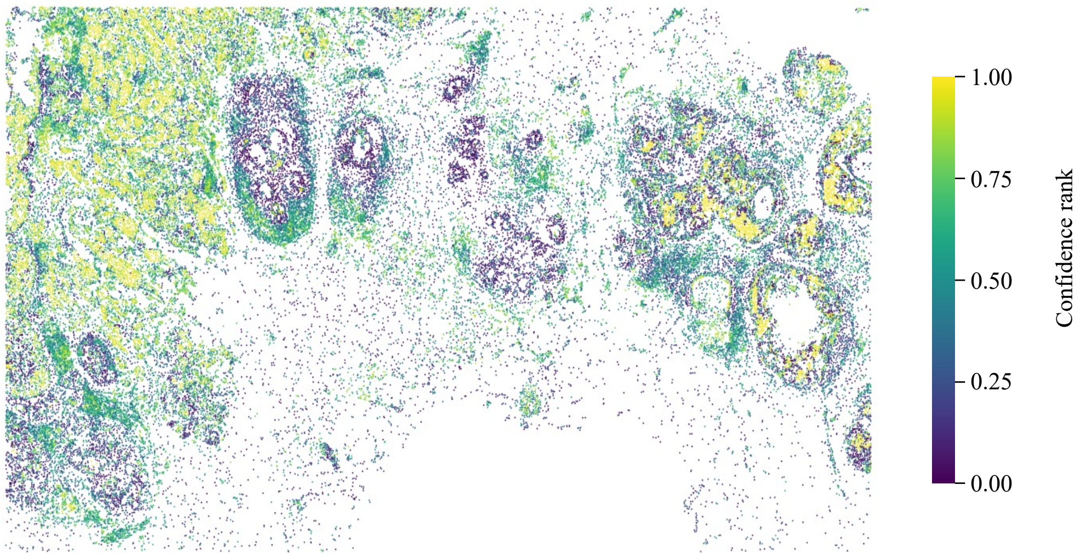


**Supplementary Fig. 3 | Spatial distribution of cell-level prediction confidence in the breast1rep2 cross-section test sample.** Spatial visualization of REGAIN prediction confidence across individual cells. Confidence scores are represented as normalized confidence ranks ranging from 0 to 1, with higher values indicating greater estimated prediction reliability.

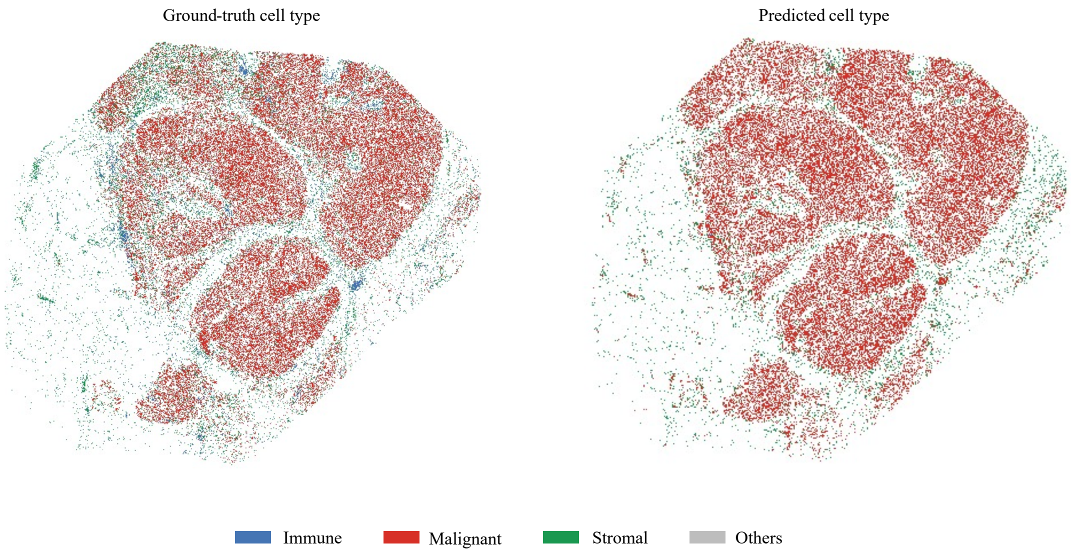


**Supplementary Fig. 4 | Comparison of ground-truth and REGAIN-predicted cell-type distributions in the cross-section breast cancer dataset.** Cell types were grouped into four categories: malignant, immune, stromal and other cells. Ground-truth cell-type annotations are shown on the left, and REGAIN-predicted cell types are shown on the right.

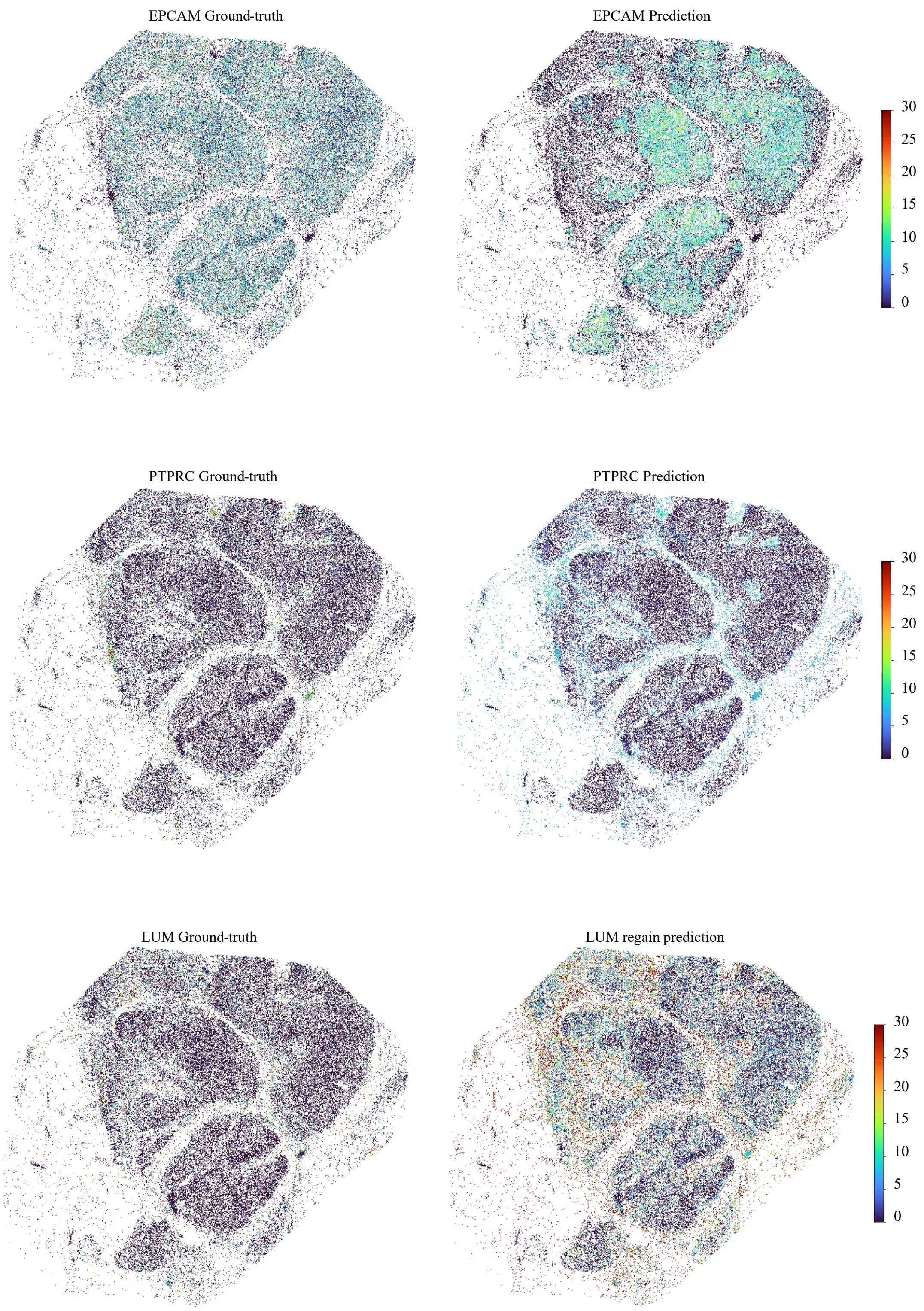


**Supplementary Fig. 5 | Spatial visualization of representative marker genes in the TENX94 cross-dataset test sample.** Spatial expression patterns of EPCAM (malignant epithelial/tumor-cell marker), PTPRC (immune-cell marker) and LUM (stromal-cell marker). For each gene, ground-truth expression is shown in the left column and REGAIN-predicted expression is shown in the right column.

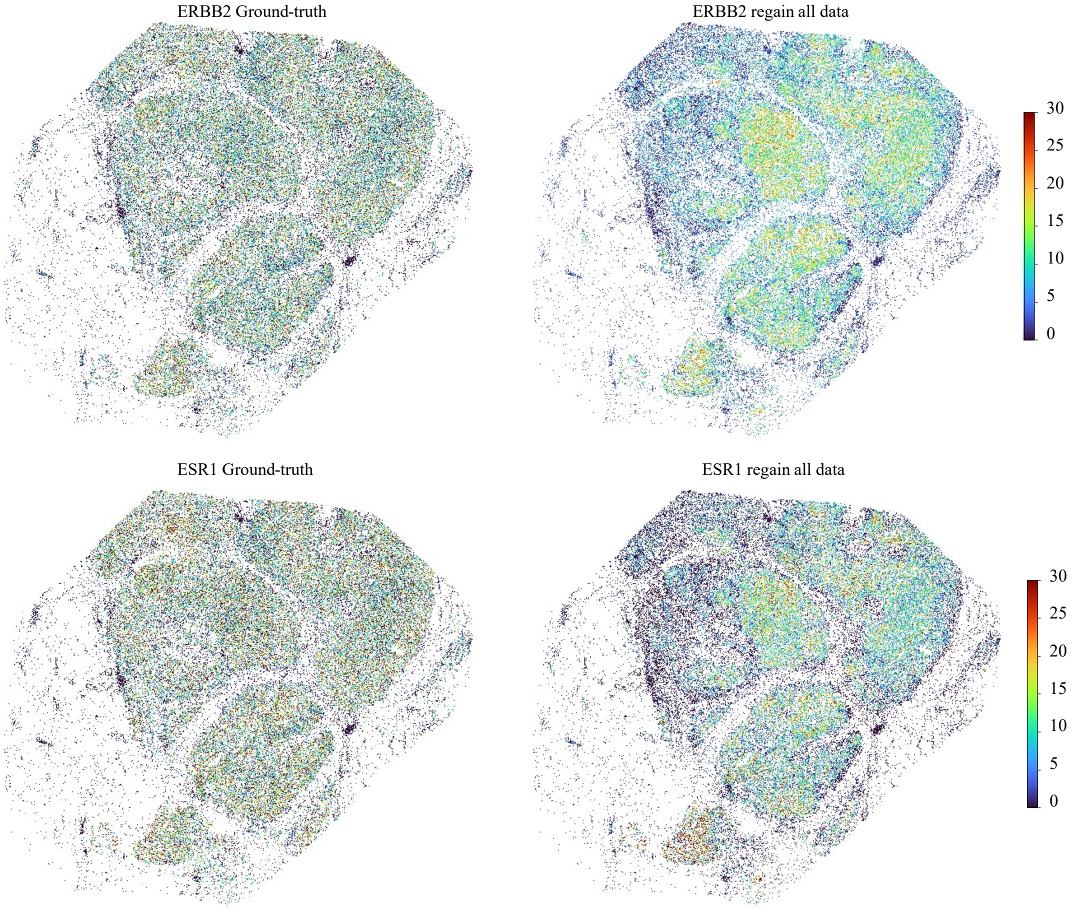


**Supplementary Fig. 6 | Spatial visualization of additional breast cancer-associated marker genes in the TENX94 cross-dataset test sample.** Spatial expression patterns of ERBB2 and ESR1. For each gene, ground-truth expression is shown in the left column and REGAIN-predicted expression is shown in the right column.

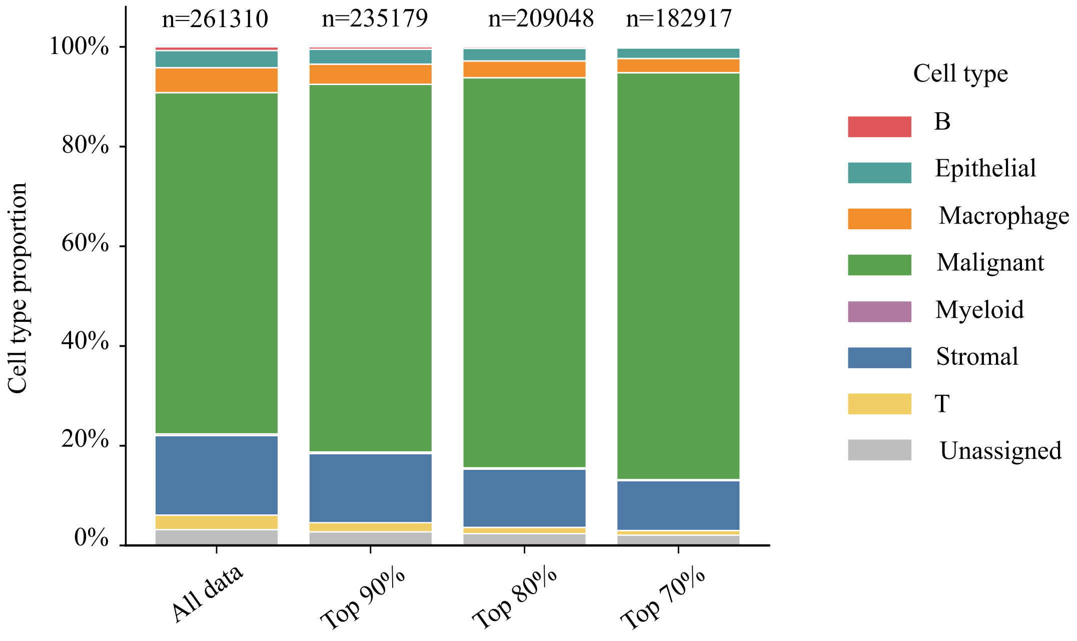


**Supplementary Fig. 7 | Cell-type composition changes under confidence-guided filtering in the TENX94 cross-dataset test sample.** Cell-type proportions across all cells and subsets retaining the top 90%, 80% and 70% most confident cells. The number of retained cells (n) is indicated above each bar.

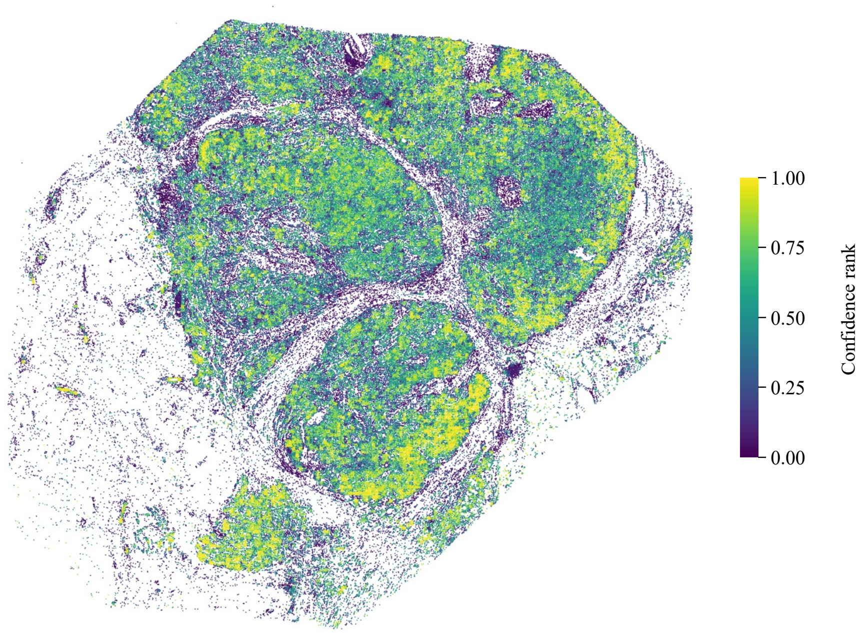


**Supplementary Fig. 8 | Spatial distribution of cell-level prediction confidence in the TENX94 cross-dataset test sample.** Spatial visualization of REGAIN prediction confidence across individual cells. Confidence scores are represented as normalized confidence ranks ranging from 0 to 1, with higher values indicating greater estimated prediction reliability.

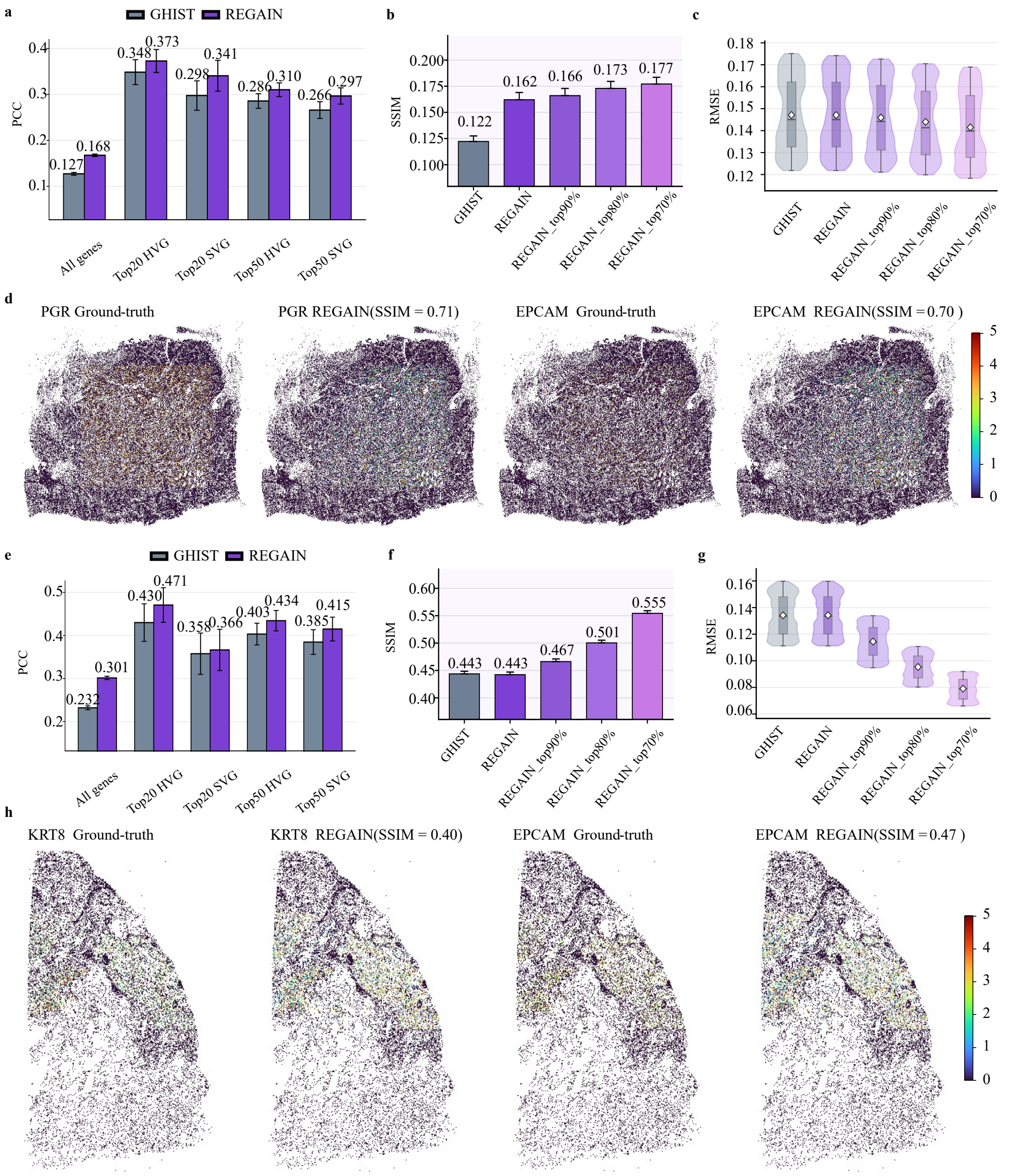


**Supplementary Fig. 9 | Xenium-to-Visium HD cross-platform transfer in breast and lung cancers.** (a) Breast cancer: gene-wise PCC comparison between REGAIN and GHIST for all genes, HVGs and SVGs. (b) Breast cancer: gene-wise SSIM comparison for GHIST, REGAIN and REGAIN predictions after confidence-guided filtering. (c) Breast cancer: cell-wise expression RMSE comparison for GHIST, REGAIN and confidence-filtered REGAIN predictions. (d) Spatial visualization of representative breast cancer marker genes, including PGR and EPCAM, showing ground-truth and REGAIN-predicted expression. (e) Lung cancer: gene-wise PCC comparison between REGAIN and GHIST for all genes, HVGs and SVGs. (f) Lung cancer: gene-wise SSIM comparison for GHIST, REGAIN and REGAIN predictions after confidence-guided filtering. (g) Lung cancer: cell-wise expression RMSE comparison for GHIST, REGAIN and confidence-filtered REGAIN predictions. (h) Spatial visualization of representative lung cancer marker genes, including KRT8 and EPCAM, showing ground-truth and REGAIN-predicted expression.

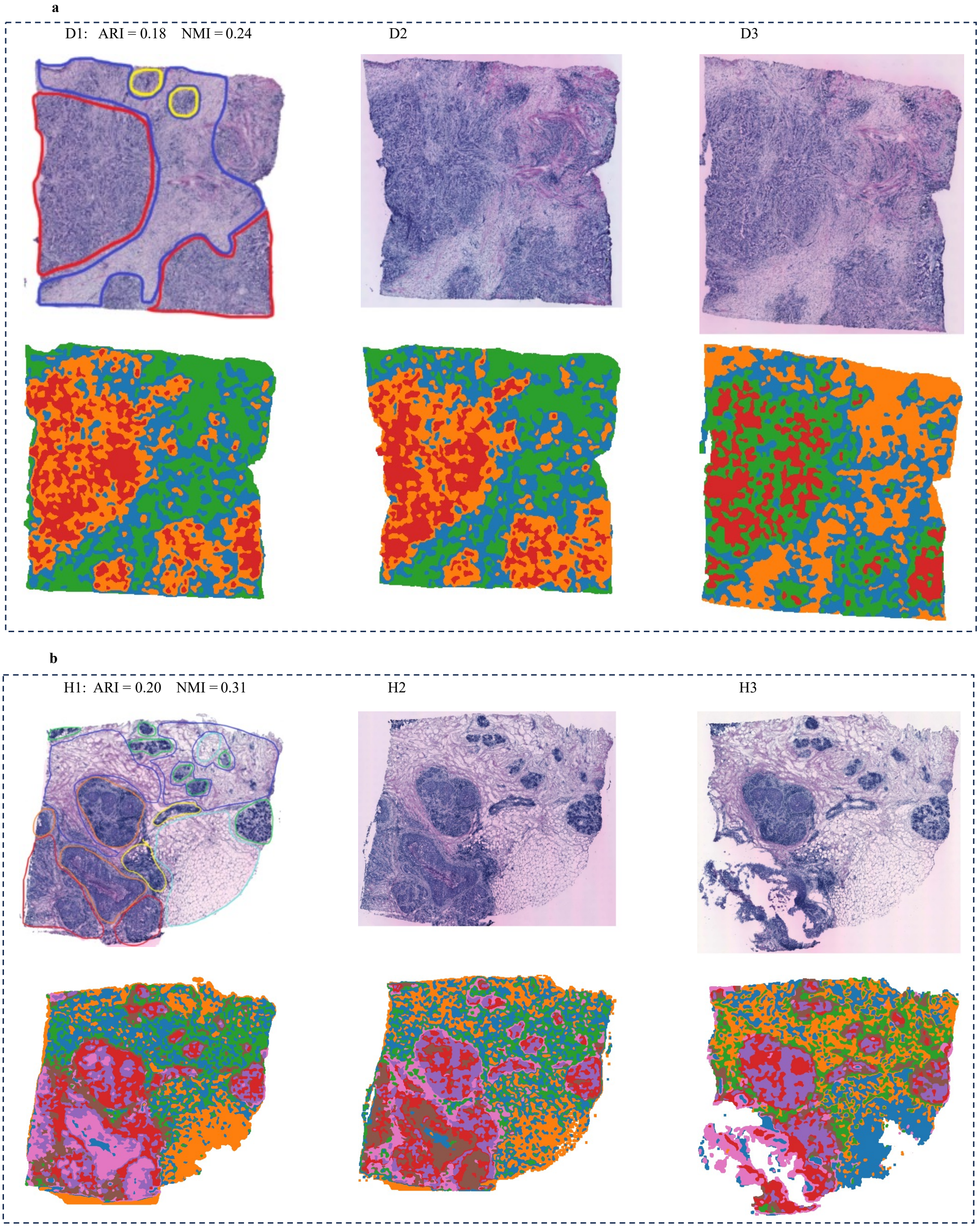


**Supplementary Fig. 10 | Additional spatial-domain results on the HER2ST spot-level dataset.** (a) H&E images and spatial-domain results for samples D1, D2 and D3. (b) H&E images and spatial-domain results for samples H1, H2 and H3. These panels complement the HER2ST spatial-domain results shown in the main figure.
